# Source-Decomposed Multimodal Graph Connectivity: A Temporal Mixture-Manifold Pipeline for Tracking Ischemic Injury and Treatment

**DOI:** 10.64898/2026.10.11.758347

**Authors:** Hyoshin Park, Jae Lee, Aditya Nimmagadda, Brian Llinas, Mandana Farhang Ghahfarokhi, Alberto Musto, Seungjun Ryu

**Affiliations:** Engineering Management & Systems Engineering, Old Dominion University, Norfolk, VA, USA; Mechanical & Aerospace Engineering, Old Dominion University, Norfolk, VA, USA; Computer Science, Old Dominion University, Norfolk, VA, USA; Biomedical & Translational Science & Neurology, Old Dominion University, Norfolk, VA, USA; Neuroscience, Johns Hopkins University, Baltimore, MD, USA

**Keywords:** dynamic functional connectivity, mixture model, graph network construction, Ensemble Kalman Filter, diffusion tensor imaging, resting-state fMRI

## Abstract

To capture the complex, dynamic network reorganization following acute focal brain injuries that standard static or unimodal neuroimaging frameworks routinely miss, we present an open, reproducible computational multimodal pipeline that integrates resting-state functional MRI and diffusion tensor imaging (DTI) into dynamic, source-decomposed graph networks. This framework preserves sub-voxel, multi-peak distributional dynamics over time by modeling regional blood-oxygen-level-dependent signals within sliding windows using a Gaussian mixture model, with model complexity dynamically optimized. The resulting pairwise similarities define time-varying edge weights across local, nonlocal, and DTI-supported structural sources, unifying them into a multimodal connectivity manifold that precisely quantifies node strength, topological turnover, and boundary modifications. Applied to a non-human primate model of photothrombotic ischemia, the methodology revealed severe post-injury disruptions, with 66 of 144 nodes (45.8%) fundamentally altering their connectivity across localized sources. Furthermore, coupling this architecture with an Ensemble Kalman Filter (EnKF) allowed for the continuous tracking of a tissue “reversibility” coordinate, yielding earlier detection of metabolic state collapse than unimodal baselines and generating a forecast-divergence index as a rigorous early-warning signal. Extended across post-injury treatment boundaries to evaluate network recovery toward baseline, this interpretable, modality-aware framework establishes a standardized, shareable methodological foundation for tracking complex pathophysiological dynamics in neuroinformatics research.

## 1 Introduction

Acute injury to the central nervous system does not progress as a discrete, synchronized transition from viable to irreversibly damaged tissue. Instead, it evolves as a spatially heterogeneous, nonstationary spatiotemporal process in which neighboring micro-domains occupy highly distinct, fluctuating metabolic states, and in which network-level topology is continuously reshaped over multiple timescales. Downstream pathophysiological cascades (e.g., including excitotoxicity, excitation–inhibition imbalances, distant diaschisis, and microvascular instability) overlap nonlinearly in space and time. This produces an evolving tissue landscape that is neither uniformly healthy nor uniformly lost. Extracting actionable insight from this nonstationary process requires neuroinformatics pipelines that provide computational representations that are simultaneously *dynamic*, capturing multi-scale reorganization over time, and *heterogeneity-preserving*, ensuring that co-located but distinct cellular or metabolic sub-populations are not collapsed into a homogenized regional average.

The high stakes of this representational gap are clearest in acute neurocritical care and longitudinal modeling, where therapeutic windows and tissue salvageability must be quantified while the underlying biological state is actively transitioning and only partially observed. In this setting, flattening a volatile, spatially mixed physiology into static, categorical summaries can mask the phase transitions that determine long-term clinical and structural outcomes. Regions that are transiently and functionally suppressed, yet structurally salvageable, may become indistinguishable from adjacent zones progressing toward irreversible metabolic collapse when spatially heterogeneous signals are reduced to coarse averaging schemes. To address this limitation, this study establishes an interpretable, multimodal neuroinformatics pipeline that explicitly retains sub-voxel, mixed biological states by pairing high temporal resolution with preserved within-region distribution profiles. We treat the longitudinal focal-ischemia recording utilized herein as a severe, natural perturbation with which to systematically demonstrate, stress-test, and validate our computational framework.

Most contemporary connectivity pipelines and brain-state analyses utilizing functional magnetic resonance imaging (fMRI) satisfy at most one of these two foundational requirements. Static functional connectivity collapses an entire recording into a single time-averaged correlation matrix, discarding the nonstationary temporal evolution that serves as the primary signal of interest following acute neurological insult (Calhoun, Miller, Pearlson, & Adalı, 2014; Hutchison et al., 2013). Sliding-window and dynamic functional connectivity (dFC) approaches restore the temporal axis (Allen et al., 2014; Preti, Bolton, & Van De Ville, 2017), but the modeling architectures that dominate the field (e.g., summarizing each region of interest (ROI) by a scalar mean regional time series or by a single-peak Gaussian amplitude distribution) strip away the sub-voxel heterogeneity that distinguishes transitioning tissue from stable tissue. Crossing fibers, mixed metabolic zones, and asynchronous electrophysiological signatures all manifest as multimodal, non-Gaussian configurations at the level of a localized voxel patch. Complex fiber configurations alone occupy the majority of white-matter voxels (Behrens, Berg, Jbabdi, Rushworth, & Woolrich, 2007; Jeurissen, Leemans, Tournier, Jones, & Sijbers, 2013), and even the BOLD amplitude distribution itself departs from Gaussianity (Hanson & Bly, 2001). Therefore, a single-peak summary cannot, by construction, represent these concurrent dual processes. Similarly, while whole-brain Hidden Markov Models (HMMs) and global Gaussian Mixture Models (GMMs) successfully recover discrete, recurring macro-scale network configurations (Ezaki, Himeno, Watanabe, & Masuda, 2021; Vidaurre, Smith, & Woolrich, 2017), they typically evaluate a single, global state probability at each temporal increment, rather than preserving localized, multi-peak density structures region by region.

Graph-theoretic and dynamic network architectures are nonetheless an exceptional fit for the neuroinformatics literature, where the scholarly emphasis falls on open-source shareable data structures, reproducible workflows, and the rigorous evaluation of how upstream methodological parameterization affects derived graph topologies. Recent work in this domain includes advanced graph representations of fMRI networks with explicit evaluations of motion sensitivity and test–retest reliability (Poudel, Sharma, Lorenzetti, Parsons, & Cerin, 2024), rigorous comparisons of empirical state-transition configurations, and personalized, cloud-deployable whole-brain network models that are now entering prospective clinical evaluation, such as the Virtual Epileptic Patient and the EPINOV trial (Jirsa et al., 2023). These efforts share a foundational premise: that the scientific utility of a derived brain network is intrinsically linked to how it is constructed, and that these algorithmic construction choices deserve to be made explicit, decomposed, and globally shared rather than buried within an opaque, static adjacency matrix.

What remains missing in the informatics landscape is a unified pipeline that: (i) treats the full *distributional* (mixture) density structure of each local region as the foundational primitive from which network connectivity is built, rather than reducing each region to a scalar average; (ii) makes explicitly clear *which specific source* of connection (i.e., local spatial neighborhood, nonlocal distributional similarity, or structurally supported pathways) drives any observed network alteration, rather than reporting an undifferentiated, fused weight; and (iii) is packaged for open-science reuse, complete with documented and deposited per time and reason specific intermediate output data structures, in alignment with modern resource-sharing expectations. Each of these components is individually computationally feasible, but the integration of a dynamic, mixture-preserving, source-decomposed, shareable graph construction has not yet been operationalized and demonstrated as a single, cohesive analysis tool.

We address this gap with an open computational framework termed the Temporal Mixture Manifold Learning with Diffusion Tensor Integration (TMML-DTI), which derives source-decomposed dynamic graph networks from co-registered rs-fMRI and DTI data. The pipeline rests on three primary architectural design decisions:

1. **Order-Adaptive Mixture Primitives:** The core connectivity primitive is a localized, nodal mixture manifold whose optimal complexity and model order (one to four distinct components) are dynamically selected using the Bayesian theory, natively preserving multi-peak sub-voxel amplitude structures within short, 30-second sliding windows.
2. **Optimal-Transport Similarity Mapping:** Pairwise edge weights between topological nodes are computed via a monotonic transform of the Wasserstein distance between their fitted continuous densities, ensuring two regions are connected based on the structural topology of their underlying *distributions* rather than mere linear correlations.
3. **Multi-Source Graph Decomposition:** The resulting dynamic network is explicitly decomposed into three interpretable edge sources (e.g., local spatial connectivity, nonlocal distributional similarity, and DTI-supported structural matrix edges) unified into a multimodal connectivity manifold. This geometry-based structural decomposition permits node- and edge-level topological shifts to be attributed to specific physical or functional mechanisms rather than to an averaged network effect.

In summary, the core methodological contributions of this paper are:

- A structural connectivity primitive based on order-adaptive nodal mixtures that natively preserves within-region multi-modality over time;
- An optimal-transport similarity framework between fitted densities that defines mixture-aware, non-parametric edge weights;
- An explicit decomposition of the dynamic graph into local, nonlocal, and structurally supported sources that remain fully separable and traceable following combination;
- A suite of additive, source-traceable graph summaries (specifically node strength, topological graph turnover, and signed boundary change metrics) optimized for longitudinal before/after comparison;
- A complete repository of open-science intermediate data structures, containerized environments, and summary outputs that render every figure within this manuscript fully regenerable and every claim computationally verifiable; and
- An online streaming module mapping a per-node reversibility coordinate via a mixture-preserving Ensemble Kalman Filter (EnKF), generating a mathematically rigorous Forecast Divergence Index (FDI) as an early-warning metric and demonstrating that the underlying mixture representations support predictive tracking fidelity on par with explicit spatial connectivity analysis.

We demonstrate the pipeline on a longitudinal before/after focal-ischemia non-human primate (NHP) recording, treating the acute injury as a severe, natural perturbation with which to evaluate network reorganization. We emphasize at the outset that this investigation serves as a methodological proof of concept and an infrastructure contribution: the derived graphs represent descriptive, modality-aware temporal summaries, demonstrating strong weighted connectivity within the constructed graph schema.

The remainder of the paper is organized as follows. Section 2 situates the method against prior work on dynamic functional connectivity, mixture modeling, optimal-transport similarity, and multimodal data fusion. Section 3 specifies the architecture including preprocessing, windowing, mixture estimation, metric tracking, graph construction, and the summary statistics used for boundary comparison. Section 4 reports the empirical validation and preclinical demonstration. Section 6 discusses the neuroinformatics contribution, its relation to a broader closed-loop estimation framework, and its limitations, and Section 7 concludes the paper.

## 2 Background And Related Work

Within the neuroinformatics literature, connectivity pipelines are best evaluated along two primary computational dimensions rather than as a flat sequence of preprocessing stages. The first dimension concerns the computational representation of regional activity; whether a localized region is reduced to a unimodal summary statistic (such as a scalar mean or a single Gaussian distribution) or preserved as a heterogeneity, retaining distribution (a mixture). The second dimension concerns the topological structure imposed on these representations; whether the pipeline tracks dynamic pairwise coupling over time (Vidaurre et al., 2017), or constructs an explicit, multimodal network, connectivity geometry where edges carry both a mathematically rigorous distance and a separable source label (Poudel et al., 2024; Su et al., 2025).

As mapped out in Table 1, existing literature heavily populates three of the four possible methodological intersections: dynamic connectivity of scalar means (Vidaurre et al., 2017), structural-functional graph fusion based on scalar weights (Jirsa et al., 2023; Poudel et al., 2024; Su et al., 2025), and global mixture-based state summaries over time (Ezaki et al., 2021). The present TMML-DTI framework addresses the remaining computational space, pairing a regional, mixture-based primitive with an explicit network-connectivity geometry to keep structural and functional sources mathematically separable.

**Table 1.** Literature and contributions of the TMML-DTI architecture.

| Concepts | Prior Paradigms | Representational Gap | This Study (TMML) |
| --- | --- | --- | --- |
| **Dynamic Coupling** | Sliding-window correlations of *scalar means*; latent HMM states (Vidaurre et al., 2017). | Scalar means mathematically erase sub-voxel heterogeneity; network parameters are often opaque and unshared. | Replaces scalar summaries with an explicit, **localized distribution primitive** (Park et al., 2022), adopting strict open-science reproducibility norms. |
| **Signal Representation** | Global GMM/HMM state summaries (Ezaki et al., 2021). | Global models lose localized biological variance; unimodal nodes hide mixed, transitioning tissue states. | Fits an **order-adaptive spatiotemporal mixture**, natively preserving multimodality to identify compromised vs. recoverable tissue. |
| **Network Geometry** | Structural–functional graph fusion based on standard *scalar* edge weights (Jirsa et al., 2023; Su et al., 2025). | Standard distance metrics fail on complex distributions; premature matrix fusion prevents modality attribution. | Derives edges via **optimal-transport** distances on a **candidate DTI scaffold**, keeping structural and functional sources mathematically separable. |

### 2.1 From Static To Dynamic Functional Connectivity

Functional connectivity, classically estimated as the Pearson correlation between regional fMRI time series over an entire scan, has been productive, but it assumes stationarity: a single number is taken to describe the relationship between two regions for the whole recording. Dynamic functional connectivity relaxes this assumption, most commonly through sliding windows in which connectivity is re-estimated within short, overlapping segments, or through state-based models that assign each time point to one of a few recurring connectivity configurations (Vidaurre et al., 2017). These approaches have revealed that between-region coupling fluctuates on the scale of tens of seconds and that the statistics of these fluctuations carry information about cognition and disease (Vidaurre et al., 2017).

However, derived dynamic networks are sensitive to construction choices such as window length, thresholding, and the summary statistic used per region and that this sensitivity is not always reported. The literature has responded by foregrounding reliability and reproducibility of network construction (Poudel et al., 2024), a challenge that this paper is overcoming by decomposing and releasing every graph source. Three techniques of dynamic-connectivity construction have been commonly used:

1. **Re-estimating pairwise statistical dependence within sliding windows** typically by windowed correlation or its regularized variants, and then summarizing the resulting sequence of matrices by i) clustering windows into recurring connectivity states, ii) tracking graph-theoretic indices over time, or iii) computing the variability of each edge (Su et al., 2025).
2. **Dispensing with explicit windows and instead inferring latent states directly**, as in hidden Markov models that assign each volume to one of a few configurations (Vidaurre et al., 2017), or in co-activation-pattern analyses that cluster individual time points by their spatial activity maps.
3. **Preserving heterogeneity** by retaining short windows but changing the *primitive* from which connectivity is computed (Park et al., 2022).

Instead of a windowed correlation of mean signals, this framework derives an explicit perregion descriptor, a fitted amplitude distribution, and connects regions by comparing those descriptors (Park et al., 2022). This paper takes advantages of interpretability of the primitive; its cost is the need to estimate a richer per-region quantity, which is why window length and model-order control matter. Across all three category of connectivity literature, the field increasingly recognizes that the *construction recipe* is part of the result, and that releasing intermediate products is necessary for others to reproduce and re-interpret a derived network (Poudel et al., 2024).

### 2.2 Mixture Models Of Regional Signal Distributions

A complementary line of work models the *distribution* of regional activity rather than its time-ordered trajectory. GMMs and HMMs can summarize resting-state signals into a small number of states (Ezaki et al., 2021); the comparison between them shows that the mixture model, which ignores temporal ordering, performs comparably to the hidden Markov model when records are short or temporal resolution is low, and is attractive when the goal is to characterize the instantaneous distribution of activity rather than the transition dynamics (Ezaki et al., 2021). Our use of mixtures differs in granularity and purpose (Park et al., 2022): rather than fitting a small number of global states, we fit a mixture *per node and per window* and retain its order (*k*) and shape as a local descriptor of heterogeneity (Park et al., 2022). The selected order itself becomes an interpretable quantity (e.g., higher *k* indicating a more multi-modal local amplitude distribution) and its change over time is a primary readout (Section 4.1).

The motivation for preserving multi-modality is physiological as well as statistical. Within a small patch of tissue, the measured signal aggregates contributions from sub-populations that need not behave identically: distinct metabolic or neuronal compartments, partial-volume mixtures at tissue boundaries (Friston, Harrison, & Penny, 2003), and after injury; coexisting recoverable and compromised micro-environments. When such sub-populations contribute amplitude values drawn from different ranges, the aggregate within-window distribution is genuinely multi-modal, and collapsing it to a single mean or a single Gaussian discards the structure that signals heterogeneity. A mixture representation makes this heterogeneity explicit and quantifiable: the number of components is a coarse index of how many distinguishable amplitude regimes are present, while the component means, variances, and weights describe their locations and relative prevalence. Selecting the order by a penalized criterion such as BIC, rather than fixing it, ensures that additional components are admitted only when the data support them, so that a node reported as multi-modal is not merely overfit. This local, order-adaptive description is the feature that the rest of the pipeline builds upon: similarity, graph construction, and the before/after comparison all operate on these fitted distributions rather than on scalar summaries (Park et al., 2022).

### 2.3 Optimal-Transport Similarity Between Distributions

Comparing two fitted mixtures requires a distance between probability distributions. The Wasserstein (optimal-transport) distance is a natural choice because it respects the geometry of the underlying value space: it measures the minimal cost of transporting probability mass from one distribution to the other and remains meaningful when distributions have different numbers of modes or little overlap, situations in which divergences such as Kullback–Leibler are unstable or undefined (Delon & Desolneux, 2020). Converting this distance into a bounded similarity via a decaying exponential (Section 3.2.2) yields edge weights that are large when two nodes share distributional shape and small when they do not, providing a principled, mixture-aware alternative to correlation of mean signals.

Several properties make the optimal-transport distance well suited to comparing fitted mixtures specifically (Chen, Georgiou, & Tannenbaum, 2019). It is a true metric on the space of probability distributions, so the induced similarities behave coherently: nodes that are each similar to a third node are constrained in how dissimilar they can be to one another. It is sensitive to the *location* of probability mass, so two distributions that differ by a shift are judged different in proportion to the shift, whereas overlap-based measures saturate. And it degrades gracefully when two distributions have disjoint support, returning a large but finite distance rather than the divergent or undefined values that information-theoretic measures can produce in the same situation (Chen et al., 2019; Delon & Desolneux, 2020).

These are the regimes that arise when comparing a unimodal node to a tri- or quadri-modal node, or comparing nodes before and after a perturbation that shifts amplitude regimes. For one-dimensional distributions the transport distance is inexpensive to evaluate, which matters because the pipeline computes pairwise distances over 144 nodes within every one of the 44 analysis windows; the per-window normalization described in Section 3.2.2 then places the resulting weights on a comparable scale across time.

### 2.4 Structural Support And Multimodal Fusion

Functional similarity alone does not indicate whether two regions are anatomically capable of direct communication. DTI provides complementary structural evidence through estimated white-matter connectivity (Su et al., 2025). Rather than treating structural and functional graphs as separate products, this paper uses DTI edges as structural *candidates* that are then reweighted by functional (distributional) similarity, so that a structural connection contributes to the dynamic graph in proportion to the current functional agreement of the regions it links. This is deliberately conservative: the DTI source never asserts dynamic coupling on its own, and its dynamic weight is therefore not a measure of pure tract strength. Keeping the three sources separate, and only summing them into a combined graph as a final step, preserves the ability to ask which modality is responsible for any observed change; an ability that is lost when fusion happens before analysis (Park et al., 2022). Keeping the sources separate is the design choice that most distinguishes this construction from a conventional fused connectivity matrix, and it is what makes the attribution analyses in Section 4 possible.

The relationship between structural and functional connectivity is itself an active research topic: white-matter architecture constrains, but does not fully determine, the functional coupling observed at rest, and the correspondence between the two varies across regions and changes with state and pathology. A construction that fuses structure and function into a single number commits, in advance, to one view of how they relate. By contrast, the candidate-and-reweight scheme used here keeps the structural prior explicit and lets the functional similarity modulate it window by window, so that the contribution of an anatomically plausible edge rises and falls with the current functional agreement of its endpoints. This is conservative in two senses: a structural edge contributes nothing in a window where its endpoints are functionally mismatched, and the functional similarity can never invent a structural connection that the DTI did not propose. The resulting structural source is therefore a function-modulated anatomical scaffold rather than an independent assertion of coupling, and the combined graph (Eq. 6) inherits this interpretability because the share contributed by each source remains fully recoverable.

### 2.5 Network Propagation And The Spread Of Injury Effects

A further reason to retain both nonlocal and structural edges is that the effects of a focal insult are not confined to the lesion. Remote functional changes in regions structurally connected to, but spatially distant from, the injury (the phenomenon of diaschisis) are well documented, and network models of injury routinely find that disruption propagates along connectional pathways rather than diffusing isotropically through space (Jirsa et al., 2023). Graph-based propagation operators, in which a regional quantity is spread to its connectome neighbours, are a common way to model this spread (Jirsa et al., 2023). Whole-brain network models and criticality-based accounts of disordered consciousness similarly establish that recovery and collapse are network-level phenomena (Rocha et al., 2022).

This paper does not fit such a predictive propagation model; instead, it provides the substrate on which propagation could later be examined, by making both the nonlocal distributional links and the structurally supported links explicit and separable. If injury effects spread preferentially along structurally supported routes, that should appear as boundary change concentrated in the DTI source; if instead distant regions reorganize together without direct structural support, that should appear in the nonlocal source. Keeping the sources separate is therefore not only a matter of interpretability but a prerequisite for asking, in future work, *how* reorganization spreads.

### 2.6 Synthesis And Neuroinformatics Positioning

Against this background, the TMML-DTI pipeline is best understood not as a new state model or a new tractography method, but as a *construction and decomposition tool* (Deshpande & Park, 2024; Ghahfarokhi, Park, Pandey, & Yoon, 2026; Ghavidel, Park, Kovacic, & Souza-Poza, 2026; Javadnejad, Park, Kovacic, & Sousa-Poza, 2025; Park et al., 2022): it specifies how to turn co-registered fMRI and DTI into a dynamic, mixture-preserving, source-labelled graph, and how to summarize that graph for before/after comparison around an event of interest. Framed this way, the appropriate criteria for evaluating it are those one applies to a tool: whether each step is well specified, whether its parameters and their effects are documented, whether its outputs are released in a usable form, and whether it can be applied to data beyond the single example shown. We have tried to meet these criteria explicitly, and we regard the single-recording demonstration as illustrating the tool rather than validating a scientific hypothesis.

The broader research programme from which it is drawn aims at closed-loop, uncertainty-aware state estimation for neurocritical decision support (Alla, Bheesetty, & Park, 2025; Darko et al., 2022; Folsom, Ono, Otsu, & Park, 2021; Folsom, Park, & Pandey, 2022; Park, Haghani, Gao, Knodler, & Samuel, 2018); that programme, including ensemble filtering and forecast-divergence diagnostics (Evensen, 2009), is outside the scope of this paper and is referenced only to explain provenance. Here we isolate and document the data-analysis component so that it can be evaluated, reused, and extended on its own terms. The emphasis on releasing intermediate products and documenting construction choices is deliberately aligned with the resource-sharing norms of the neuroinformatics literature (Poudel et al., 2024), where a derived network is considered reproducible only when the recipe and its inputs are available to others. Complex pipelines are most useful to a community when their components can be adopted independently, and most trustworthy when each component can be validated without committing to the whole. By presenting the construction separately from the estimation machinery it was designed to feed, we make it possible for a reader to use the source-decomposed graph for a different purpose such as descriptive comparison, feature extraction, or input to another model without inheriting assumptions from the downstream programme. The design choices are not mistaken for arbitrary ones, but the contribution stands on its own as an analysis tool.

## 3 Materials And Methods

### 3.1 Experimental Paradigm And Data Acquisition

The data comprise resting-state fMRI with co-registered DTI from a NHP photothrombotic (PT) ischemic-stroke model in cynomolgus macaques (*Macaca fascicularis*). The animals, imaging hardware, injury model, and small-molecule intervention are those of the companion NHP study of Lee et al. (2025), which characterizes the same paradigm histologically, structurally, and behaviourally and this study adds the dynamic functional-connectivity analysis. The cohort comprised three macaques aged 3–4 years (2.6–5.1 kg; two male, one female), one assigned to each of three conditions: PT only (vehicle), PT followed by KDS12025 at 0.1 mg/kg/day, and PT followed by KDS12025 at 0.03 mg/kg/day.

Focal ischemia was induced by photothrombosis targeting the dorsal premotor (PMd) and primary motor (M1) cortices; the cortical territory governing hand shape and grasp under ketamine (8 mg/kg) and medetomidine (0.05 mg/kg) induction with 1.2–1.5% isoflurane maintenance: after a craniotomy exposing the motor and premotor cortices, Rose Bengal (20 mg/mL, 80 mg/kg intravenous) was activated by a 561 nm laser for 20 minutes to induce thrombosis (Lee et al., 2025). The KDS12025 intervention, a blood–brain-barrier-permeable H_2_O_2_ decomposing peroxidase enhancer that acts indirectly through haemoglobin, was administered by endotracheal intubation from approximately 10 minutes after photothrombosis, at either 0.1 mg/kg on a single day or 0.03 mg/kg once daily for three consecutive days (Lee et al., 2025). The dataset therefore spans three physiological epochs per animal (pre-injury baseline, post-injury, and post-treatment) and it is the matched pre-/post-treatment recordings that support the treatment-boundary analysis of Section 5.

Imaging was performed on a Siemens Skyra 3 T scanner (gradient insert, 45 mT/m maximum gradient, 200 mT/m/s maximum slew rate; 15-channel transmit/receive coil) (Lee et al., 2025). Resting-state fMRI was acquired with a repetition time of TR = 1.5 s (Table 2).

**Table 2.**
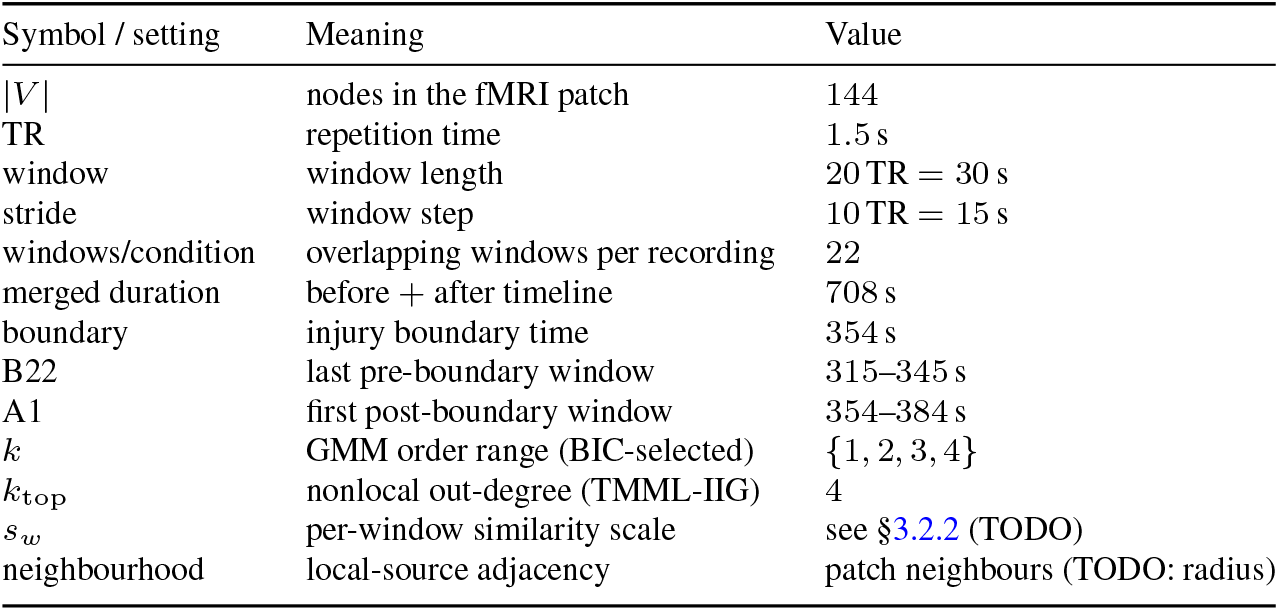
Analysis parameters and the values used in the demonstration. Values marked TODO depend on acquisition details still to be confirmed.

Co-registered DTI provided the structural candidate edges, with infarct extent quantified in 3D Slicer (Fedorov et al., 2012) and white-matter tractography; including the corticospinal tract used for the structural validation of Section 5.2 reconstructed by generalized *q*-sampling imaging in DSI Studio (Lee et al., 2025). In the reference paradigm, structural recovery was assessed at post-operative day 14 (POD14), which anchors the nominal post-treatment imaging point.

For the connectivity demonstration, a single representative axial slice provides a two-dimensional fMRI patch of 144 cells (graph nodes). The before-injury and after-injury recordings are concatenated into one merged timeline of total duration 708 s, with the injury boundary at *t* = 354 s; the last pre-boundary analysis window is denoted B22 (merged *t* = 315–345 s) and the first post-boundary window A1 (merged *t* = 354–384 s), and these two windows anchor all before/after-injury comparisons below. The matched post-treatment recording is windowed identically and contributes a parallel treatment boundary (Section 5). The dataset is deliberately compact; a controlled perturbation suitable for demonstrating and stress-testing the construction.

To ensure computational tractability and maximize the visual clarity of the network topologies, the pipeline is evaluated here using a single co-registered slice framed as a localized patch. This patch comprises a contiguous grid of nodes wherein the local source possesses a well-defined spatial neighborhood. Critically, these identical nodes map both the functional (BOLD) features utilized for mixture estimation and the structural (DTI) candidate edges, ensuring that all three graph sources are evaluated on a unified node set. Consequently, spatial co-registration of the functional and structural modalities is a strict prerequisite; this framework assumes both data types are aligned to a common stereotaxic space prior to patch extraction. Importantly, no mathematical or architectural components of this framework depend on the two-dimensionality of the patch; this restriction is a deliberate choice for illustrative clarity, with the full volumetric extension detailed in Section 6.6.

#### Temporal modeling

Each condition (before, after) is divided into 22 overlapping temporal windows. With a repetition time of TR = 1.5 s, each window spans 20 TRs (30 s) and successive windows are offset by a stride of 10 TRs (15 s), giving 50% overlap. Before-injury windows span merged time 0–345 s and after-injury windows span 354–699 s. The window length is a compromise between temporal specificity and the sample size required to fit a stable nodal mixture; the 50% overlap yields a smooth temporal readout without inflating the effective number of independent windows. The merged before/after timeline is treated as a single sequence of 44 windows so that the same descriptors and summaries apply uniformly on both sides of the boundary; the boundary itself is not modeled as a special window but is located between windows B22 and A1, which keeps the before/after contrast symmetric and free of any window that straddles the event.

### 3.2 The Dynamic Network Construction

We model the transformation of two co-registered recordings (before and after the event of interest) into a sequence of source-decomposed dynamic graphs and a set of before/after summaries. At a high level it proceeds in nine steps: (1) split each recording into overlapping temporal windows; (2) for each node and window, fit a Gaussian mixture to the windowed BOLD values; (3) select the mixture order by BIC; (4) track how the selected order changes across nodes and windows; (5) compute pairwise Wasserstein distances between fitted node densities within each window; (6) convert distances to bounded similarity weights; (7) build the three source-specific graphs (local, nonlocal top-*k*, DTI-supported) from those weights; (8) sum them into a combined graph while retaining source labels; and (9) summarize edge weights, node strengths, temporal turnover, and the before/after boundary change. Steps (2)–(6) operate purely on the functional amplitude distributions; the structural modality enters only at step (7), as a set of candidate edges to be reweighted. The remainder of this section specifies each step. Table 2 collects the analysis parameters and the values used for the demonstration.

Four architectural principles govern this pipeline, prioritizing mathematical transparency and physical interpretability over black-box optimization. First, the connectivity primitive is treated as an explicit, inspectable nodal distribution rather than a latent feature learned end-to-end; while this lacks the unconstrained flexibility of deep neural embeddings, it ensures that every statistical parameter can be directly verified and reasoned about biologically. Second, distribution similarity is evaluated using an optimal-transport metric that respects the geometry of the signal space, defining regional alignment by matching shape and location rather than simple temporal correlation. Third, the graph is decomposed into separate, explicitly labeled sources that combine via a parameter-free linear sum, preventing the loss of modality attribution common in conventional matrix fusion. Finally, downstream network diagnostics such as node strength, edge turnover, and signed boundary changes are kept additive and fully traceable to their originating sources. Ultimately, this framework intentionally trades unconstrained predictive flexibility for auditability and modular reuse, establishing a traceable toolkit for open-science replication rather than an architecture over-engineered for a single benchmark dataset.

#### 3.2.1 Nodal Mixture Estimation (GMM)

For each node *i* and each window *w*, the BOLD values within the window are modeled as Gaussian mixture:

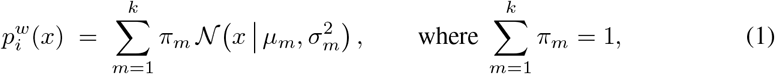

where 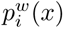 is the estimated BOLD-amplitude density for ROI (or voxel patch) *i* within temporal window *w*; *k* denotes the number of mixture components; *π_m_*, *µ_m_*, and 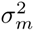 are the weight, mean, and variance of component *m*, respectively; 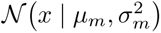 the univariate Gaussian density; and *x* is the (scalar) BOLD amplitude. The constraint on the right hand side with *π_m_ ≥* 0 makes the mixing coefficients 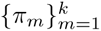 a valid probability distribution over the *k* components. The order *k* is dynamically selected from the candidate set *{*1, 2, 3, 4*}* by minimizing the BIC:

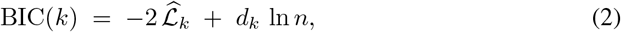

with 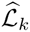 representing the maximized log-likelihood of the *k*-component fit, *d_k_* the number of free parameters, and *n* the number of temporal samples within the window. BIC explicitly penalizes parameter complexity and thus guards against spuriously high model orders in short, non-stationary windows.

The selected order 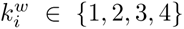 (unimodal, bimodal, trimodal, quadrimodal) is retained as a localized descriptor of tissue signal complexity, and its temporal changes are tracked across the merged timeline.

##### Stability and identifiability

Two practical concerns attend per-window mixture fitting. The first is estimation stability: with a limited number of samples per window, high-order fits can be unstable, which is precisely what the BIC penalty mitigates by demanding that the likelihood gain from an additional component outweigh its parameter cost. The selected order is therefore conservative, and a node reported as multi-modal in a window has passed this penalized test rather than merely admitting a better unpenalized fit.

The second concern is label identifiability: the components of a mixture have no inherent order, so component-wise quantities (which component is “first”) are not comparable across nodes or windows. The pipeline avoids relying on component labels by using only label-invariant quantities downstream; the selected order *k* and the Wasserstein distance between whole densities, both of which are invariant to how the components are indexed. This keeps the analysis robust to the inherent non-identifiability of mixture components.

#### 3.2.2 Optimal-Transport Distributional Similarity

Connectivity between two nodes in a window is defined directly from the topological similarity of their fitted densities. For nodes *i* and *j*, we compute the Wasserstein distance 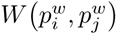 between their window-*w* mixtures and convert it into a bounded similarity edge weight. For two one-dimensional densities *F_p_* and *q* with cumulative distribution functions *F_p_* and *F_q_*, the *r*-Wasserstein distance (Peyré & Cuturi, 2019; Villani, 2009) is

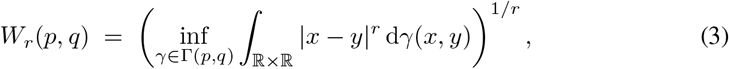

where Γ(*p, q*) is the set of all joint distributions (couplings) with marginals *p* and *q*, and *r ≥* 1 (we use *r* = 1). Intuitively, *W_r_* is the minimum “cost” of transporting the probability mass of *p* into *q*, with cost *|x − y|^r^* for moving unit mass from *x* to *y*. For univariate distributions, this admits the closed form via quantile (inverse-cumulative distribution) functions (Kolouri, Park, Thorpe, Slepčev, & Rohde, 2017),

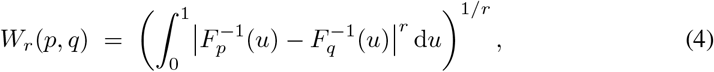

which we evaluate for each node pair to obtain 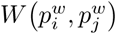 in Eq. (5).

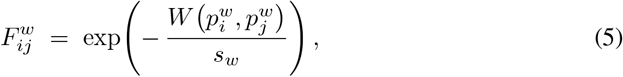

where *s_w_*is a window-specific normalization scale that balances distances within each window so that edge weights remain comparable across time.

Equation (5) yields 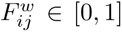, with values approaching unity when two nodes share identical distributional profiles and near zero when their distributions diverge. Because *F* is built from full fitted densities rather than from mean signals, two nodes can be judged highly similar even when their mean amplitudes differ, provided the underlying *shape* of their amplitude distributions agrees.

The per-window normalization by *s_w_*serves a specific purpose. Absolute Wasserstein distances depend on the amplitude scale of the signals within a window, which can drift over the recording; without normalization, edge weights would conflate genuine changes in distributional agreement with global changes in signal scale. Dividing by a window-specific scale places the distances on a comparable footing across windows, so that a change in *F* between B22 and A1 can be read as a change in relative distributional agreement rather than as an artefact of differing signal magnitudes. The choice of *s_w_* is consequential for the absolute magnitudes of edge weights and therefore for node strengths, and it is reported explicitly; comparisons across windows assume the same definition of *s_w_*is applied throughout. We compute the similarity field densely over all node pairs in a window and then let each source select the subset of edges it retains, so that the local, nonlocal, and structural graphs are all derived from the same underlying distance computation.

#### 3.2.3 Source-Decomposed Graph Fusion with TMML

The similarity field *F^w^* is exploited to construct three interpretable edge sources per window, which are subsequently combined. We denote the node set as *V* (*|V |* = 144).

##### Local connectivity through direct information gain (TMML-DIG)

Edges are placed strictly between spatially neighbouring nodes in the extracted patch and weighted via Eq. (5). This source captures localized distributional coordination, highlighting how tightly a node’s amplitude distribution matches its immediate anatomical surround.

##### Nonlocal connectivity through indirect information gain (TMML-IIG)

For each individual node, edges are placed to its *k*_top_ = 4 most distribution-similar *non-neighbouring* nodes, weighted by Eq. (5). This source allows long-range functional clusters to be linked when their amplitude profiles mirror each other, capturing non-contiguous coordination that a purely localized model would miss. The choice *k*_top_ = 4 fixes the out-degree of the nonlocal source; because it directly controls graph density, it is treated as an explicit tunable parameter.

##### Structural source (DTI-supported)

Co-registered DTI structural edges serve as anatomical candidates that are dynamically reweighted by the functional similarity of the nodes they connect via Eq. (5). A structural edge therefore contributes to the graph only in proportion to the current functional agreement of its endpoints; its dynamic weight is not a measure of pure tract strength.

##### Combined graph

The combined TMML-DTI graph is evaluated as a parameter-free linear sum of the three distinct source matrices:

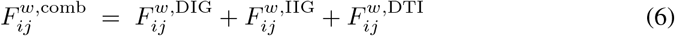

retaining, for every edge, a discrete label bitmask of which sources contributed so that overlapping (multi-source) edges remain entirely recoverable.

The three sources answer deliberately different questions and therefore occupy different parts of the graph. The local source describes how each node relates to its immediate spatial surround and so emphasizes contiguous, neighbourhood-scale coordination; in matrix form its edges fall near the diagonal once nodes are indexed by spatial proximity. The nonlocal source is, by construction, free of the spatial-proximity constraint and so populates the off-diagonal, linking distant nodes whose amplitude distributions happen to match; it is the source through which long-range coordination, or its disruption, becomes visible. The sparse structural source exists only where the DTI proposes an anatomical connection and its contribution is gated by current functional similarity.

Keeping these contributions additive and labelled means that any property of the combined graph, including a node’s combined strength or a change in it across the boundary, can be traced back to the sources responsible. The combination is intentionally a simple sum rather than a learned weighting: introducing tunable source weights would improve fit to a particular dataset at the cost of the transparency that motivates the decomposition, so we keep the combination parameter-free and report the sources separately wherever a combined quantity is shown.

### 3.3 Network Diagnostics And Topology Summaries

From each per-window graph, we extract the following diagnostics:

#### Node strength

The weighted connection strength of node *i* is defined as:

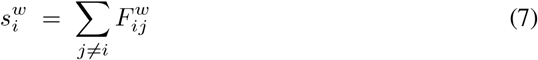

computed independently per source and for the combined topology. Node strength acts as the primary node-level readout for spatial mapping and tracking.

#### Graph turnover

Between consecutive windows, network reorganization is summarized by the mean absolute change in edge weights, treating dynamic nonlocal edges as binary indicators (present or absent per window) so that the rapid appearance and disappearance of nonlocal links contribute explicitly to the total turnover metric.

#### Boundary change

The before/after injury comparison directly contrasts window B22 with window A1. For every node and edge, we report the after-minus-before difference in strength and edge weight, signed so that increases and decreases are explicitly distinguished:

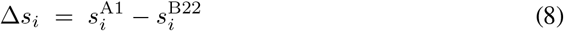

We use the single last-pre-boundary and first-post-boundary windows, rather than averages over several windows on each side, so that the contrast reflects the change closest in time to the boundary; this maximizes temporal specificity at the cost of using less data per side, a trade-off appropriate when the question is what happens *at* the boundary rather than what differs between the two conditions on average. The same boundary contrast is computed for each source and for the combined graph, so that node- and edge-level change can be reported per source as well as in aggregate.

#### Ranking and partner attribution

Nodes are ranked by the absolute boundary change in combined node strength *|*Δ*s_i_|*. For the top-ranked nodes, we additionally record their strongest connected partners before and after the boundary, along with the specific source (local, nonlocal, structural, or multi-source overlap) supporting each partner edge, yielding a prioritized target table for clinical tracking.

We emphasize the use of *absolute* change for ranking: a node may rise to the top of the ranking either by gaining or by losing connectivity, and the sign is preserved separately so that the direction of change is not lost. Ranking by absolute change foregrounds the nodes most affected by the boundary regardless of direction, which is the appropriate ordering when the goal is to identify where to look rather than to assert a directional effect. The partner attribution then adds the relational context that a node-level magnitude alone omits: it records not only that a node changed but with which other nodes its connectivity changed, and through which source, so that an affected node can be understood as part of a small sub-network rather than in isolation.

#### Visualization and statistical considerations

Spatial graph figures encode node position by patch coordinate, node colour by selected mixture order, node size/transparency by strength, edge colour by source, and edge thickness/opacity by relative weight among displayed edges. For legibility, some figures display only the strongest edges or suppress edges entirely; these display thresholds (for example, showing top-ranked edges, or the *k*_top_ rule for the nonlocal source) are presentation choices and are *not* statistical significance tests. Because the demonstration is a single recording, we report descriptive quantities only and make no inferential claims; the design of a subject-level statistical model (across multiple animals or subjects, with appropriate null models for graph density) is identified as required future work in Section 6.

### 3.4 State Tracking Via Ensemble Data Assimilation (EnKF)

The connectivity construction above characterizes *where* and *through which source* the network reorganizes. A complementary, per-cell read-out asks a different question: as data arrive over time, where does a given cell sit on a continuum between its pre-injury and postinjury state, and with what confidence? We answer this with a mixture EnKF that tracks a scalar reversibility coordinate *λ ∈* [0, 1] (*λ* = 1 fully pre-injury, *λ* = 0 fully post-injury), demonstrated here on a single representative cell (NHP fMRI Cell 52).

#### Anchor mixtures

For each cell, we fit two Gaussian mixtures once, from the 256 pre-injury and the 256 postinjury BOLD values respectively, with order chosen by BIC (*K* = 2 is well conditioned at this sample size). These two baseline densities, 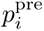 and 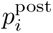, serve as the mathematical anchors for the trajectory.

#### State distribution as displacement interpolation

The reversibility coordinate indexes a family of distributions between the anchors. Rather than a linear mixture of densities, 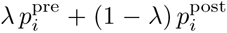 which would describe a cell *switching b*etween fully healthy and fully injured states, we interpolate the mixture *parameters* along the optimal-transport (displacement) geodesic, so that probability mass *moves* continuously from the post-injury location toward the pre-injury location as *λ* increases (Chen et al., 2019; Delon & Desolneux, 2020). For matched components *j*, the transport trajectory is governed by:

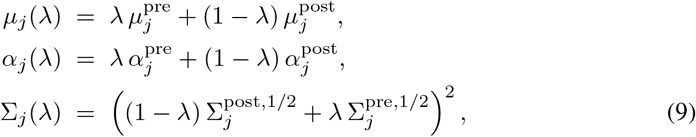

where Eq. (9) represents the Bures–Wasserstein geodesic between Gaussian covariances. The resulting time-varying structural density is expressed as:

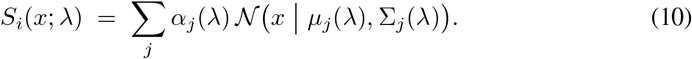

This formulation ensures that intermediate values of *λ* match intermediate biomarker profiles (partial recovery) rather than rapid binary switching, and permits *λ* to fall (secondary insult) as well as rise.

#### Ensemble filter

We now let *k* = 1, 2*, …* denote the discrete assimilation temporal index *k*. The filter maintains an ensemble of *N* members, each carrying a localized joint state vector 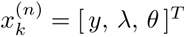 at step *k*, where (*n*) indexes the ensemble member, tracking the simulated BOLD value *y*, the reversibility coordinate *λ*, and the baseline hemodynamic parameters *θ*. The coordinate *λ* is initialized uniformly on [0, 1] to remain completely agnostic. At each time step, members are forecast through a Balloon–Windkessel hemodynamic forward model (Friston et al., 2003) with additive process noise under a standard random-walk transition:

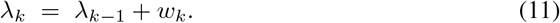

When an empirical BOLD volume *y_k_* is captured, each member’s observation likelihood is evaluated under the parameter-interpolated mixture, *L_n_* = *S_i_*(*y_k_*; *λ*^(*n*)^), and the Kalman gain updates the state vector:

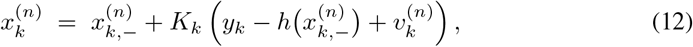

nudging every member’s latent variables toward the empirical observation (Evensen, 2009; Ullah & Schiff, 2009). We track the posterior mean E[*λ_k_*] as the current state estimate, the posterior variance Var[*λ_k_*] as the model uncertainty, and evaluate the Forecast Divergence Index (FDI):

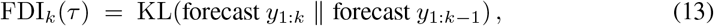

defined as the Kullback–Leibler divergence between successive *τ* -step-ahead ensemble predictions, serving as an early-warning anomaly trigger.

#### Detection comparison

To quantify the value of preserving the mixture, we compare the *K* = 2 mixture-anchor filter against a *K* = 1 (forced unimodal) filter with the identical transition model, ensemble size, gain, and update equations, so that any difference in detection timing is attributable solely to the anchor structure in the likelihood. Detection time is defined by a 2*σ* departure from the pre-injury steady-state baseline of the smoothed E[*λ*] trajectory; we also compare against a Shannon (Shannon, 1948) sliding-window variance detector. A sigmoid gate *g_k_* = 0.95 *σ* (*t −* 340)*/*8 and a posterior-driven gate *g_k_* = 1 *−* E[*λ*] illustrate how the FDI triggers an intervention window.

#### Computational implementation and complexity

The dominant computational costs are concentrated in the localized mixture estimation and the per-window pairwise optimal-transport computations. Within a single sliding window, fitting a Gaussian mixture for all *|V |* nodes across the candidate space *k ∈ {*1, 2, 3, 4*}* is strictly linear in the number of nodes and the number of samples per window (*O*(*|V |·n*)), rendering it trivially parallelizable across nodes. Evaluating the pairwise similarity field requires *O*(*|V |*^2^) onedimensional Wasserstein calculations per analysis window. With *|V |* = 144 and 44 windows, the computational footprint remains highly tractable, though the quadratic scaling in node count must be carefully monitored if expanding the patch to full whole-brain parcellations. The nonlocal source architecture compresses storage by reducing the dense *|V |*^2^ field into a sparse top-*k* format per node, while the structural source is sparse by construction. Intermediate structures are systematically cached (Section 4) to bypass redundant fitting steps during downstream figure generation.

## 4 Results

### 4.1 Temporal Evolution Of Mixture-Distribution Complexity

Across the 44 overlapping analysis windows, the BIC-selected Gaussian mixture order varied over time at all 144 imaging nodes (Figure 2). Nodes undergo an average of 14.2 *±* 3.8 model-order switches across the 44-window timeline, spending approximately 58.3% of the recording in a unimodal state (*k* = 1), 28.1% in a bimodal state (*k* = 2), 10.4% in a trimodal state (*k* = 3), and 3.2% in a quadrimodal state (*k* = 4). These values summarize model selection within the analyzed recording; because the windows overlap and no null distribution was used, they should not be interpreted as independent events or population estimates.

**Fig. 1.**
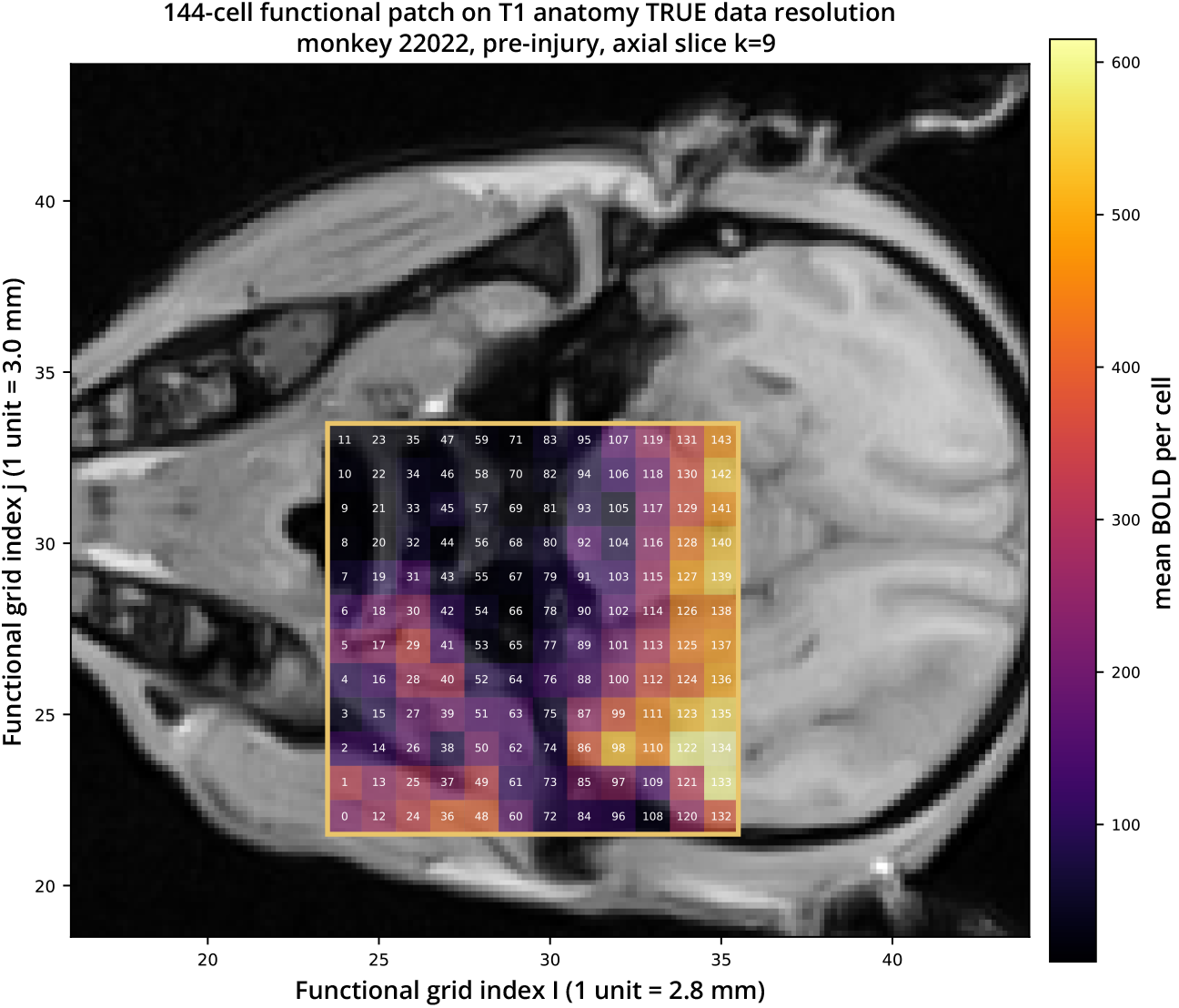
A high-resolution T1 anatomical slice (0.5 mm, greyscale) forms the background, showing an axial section through the macaque head cortex, midline, and surrounding tissue are visible. Overlaid on it is the 144-cell analysis patch, outlined by a gold box, positioned over the mid-brain. Each of the 144 cells is drawn as a single functional voxel (*∼* 2.8 *×* 3.0*mm*), a discrete square colored by its mean BOLD signal on an inferno colormap (dark purple = low, bright yellow = high), and labeled with its cell id (0-143). The numbering runs bottom-to-top within each column, then left-to-right across columns. The visible pattern: low signal (dark) on the left/upper cells, rising to high signal (yellow) in the lower-right cells (122, 133, 134). Axes are the functional grid indices i and j; a color bar on the right maps color to mean BOLD. The T1 is aligned to the functional MRI data by mutual-information cross-modal registration.

**Fig. 2.**
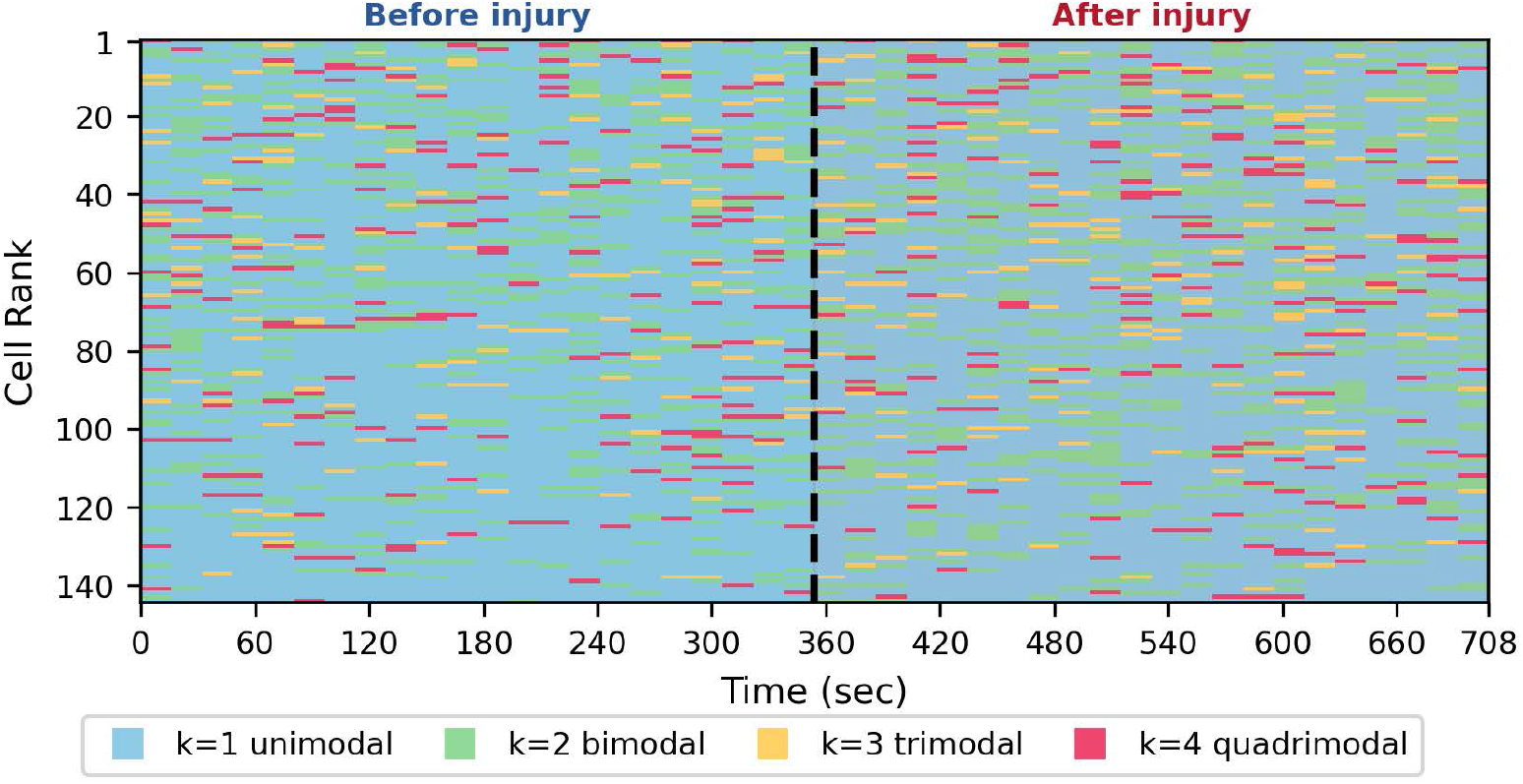
Ranked spatiotemporal evolution of cell-level multimodal distribution type (k=1-4) from NHP data **?** before and after injury, columns represent overlapping 30 s windows, and colour indicates selected GMM component count (*k* = 1 unimodal, *k* = 2 bimodal, *k* = 3 trimodal, *k* = 4 quadrimodal). The dashed vertical line marks the injury boundary at *t* = 354 s, separating the before-injury (left) and after-injury (right) segments of the merged timeline.

At the acute transition boundary, the change from the final pre-injury window (B22) to the first post-injury window (A1) was accompanied by a model-order shift in 66 of 144 nodes (45.8%). The ranked raster shows that these changes were distributed across the sampled patch rather than confined to one coordinate, with more nodes moving toward higher-order mixtures than toward (*k* = 1). This pattern is consistent with a spatially distributed change in local BOLD-amplitude distributions at the injury boundary.

Mapping selected mixture order to the native patch coordinates showed a spatially interspersed pattern of unimodal and multimodal nodes before and after the boundary. The node-by-transition raster also showed a concentration of order changes near the boundary. Together, these visualizations identify when and where the fitted distributional representation changed within this recording.

### 4.2 Connectivity Strength And Source Contributions Over Time

Figure 3 summarizes the mean edge weight and edge turnover of the source-specific and combined graphs across the concatenated timeline. Before the injury boundary, the combined mean edge weight fluctuated within a comparatively narrow range; after the boundary, both its level and variability changed. The turnover trace indicates that the identity or weight of edges also changed between adjacent windows.

**Fig. 3.**
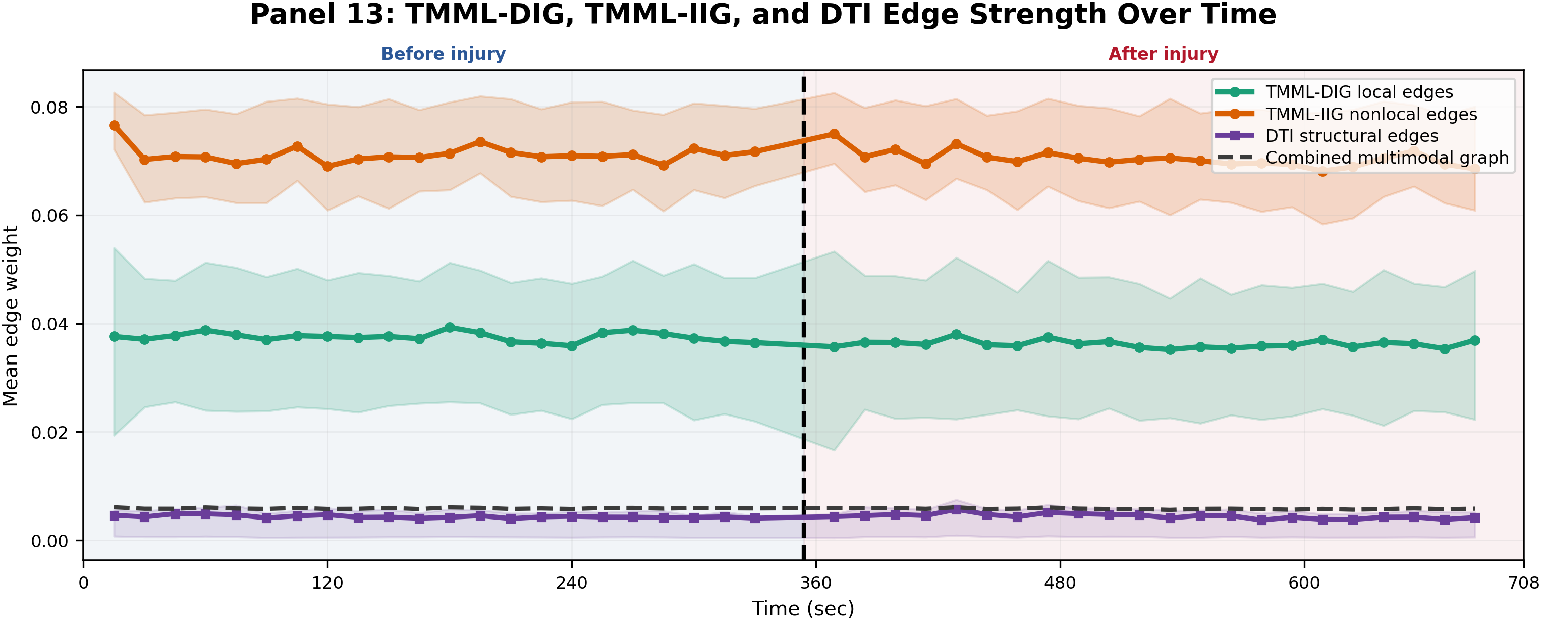
Source-wise TMML-DTI edge-strength time course over the merged timeline. Separate traces summarize TMML-DIG local neighbouring-node distributional similarity, TMML-IIG nonlocal top-*k* distributional similarity, DTI structural support reweighted by functional similarity, and the combined multimodal graph; shaded bands show the 25th–75th percentile range of source-specific edge weights. The dashed vertical line marks the injury boundary.

Across the full timeline, the nonlocal functional source (TMML-IIG) had the highest mean edge weight, the local functional source (TMML-DIG) had an intermediate mean, and the DTI-supported source had the lowest mean. This ordering is expected in part from the construction of the sources: TMML-IIG selects distributionally similar nodes across the full patch, TMML-DIG is restricted to local neighbors, and the DTI-supported graph is sparse and reweighted by time-varying functional similarity. Considering mean edge weight together with turnover separates a change in average graph weight from a change in edge configuration. At the injury boundary, both summaries departed from their pre-boundary pattern, indicating a reconfiguration of the modeled network under the prespecified graph construction.

### 4.3 Dynamic Connectivity Matrices And Spatial Layout

The adjacency matrices provide a direct view of the graph before spatial rendering. Near-diagonal entries arise primarily from the local functional source, whereas off-diagonal entries include nonlocal functional and DTI-supported edges. Off-diagonal weights were present in each representative window, and the matrix pattern changed across the injury boundary. This confirms that the spatial renderings reflect changes in the underlying edge-weight matrices rather than only changes in visualization settings.

Projecting the combined graphs into native patch space (Figure 5) provides the spatial layout of network coordination. Nodes are positioned at their patch coordinates, node size encodes total connectivity strength, node color represents selected mixture order, and edge color identifies the generating source. The representative windows show that graph changes occurred at multiple sampled coordinates in both conditions. The display is intended to localize model-derived changes within the patch.

At the first post-boundary window (A1), separating the graph into local, nonlocal, and DTI-supported sources showed that the nonlocal source contributed the largest mean edge weights under the selected top-k rule, whereas the local and DTI-supported sources were sparser by design (Figure 4). A given combined node strength can therefore arise from different mixtures of local, nonlocal, and DTI-supported edges. Source decomposition identifies which modeled source contributes to a node-level summary without assigning a causal mechanism to that contribution. A node-only rendering of the boundary window, where edge clutter is suppressed and nodes are sized strictly by total strength, highlights primary network hubs at the exact moment of injury, while the accompanying edge-resolved rendering (Figure 6) maps these hubs to their supporting functional and structural pathways.

**Fig. 4.**
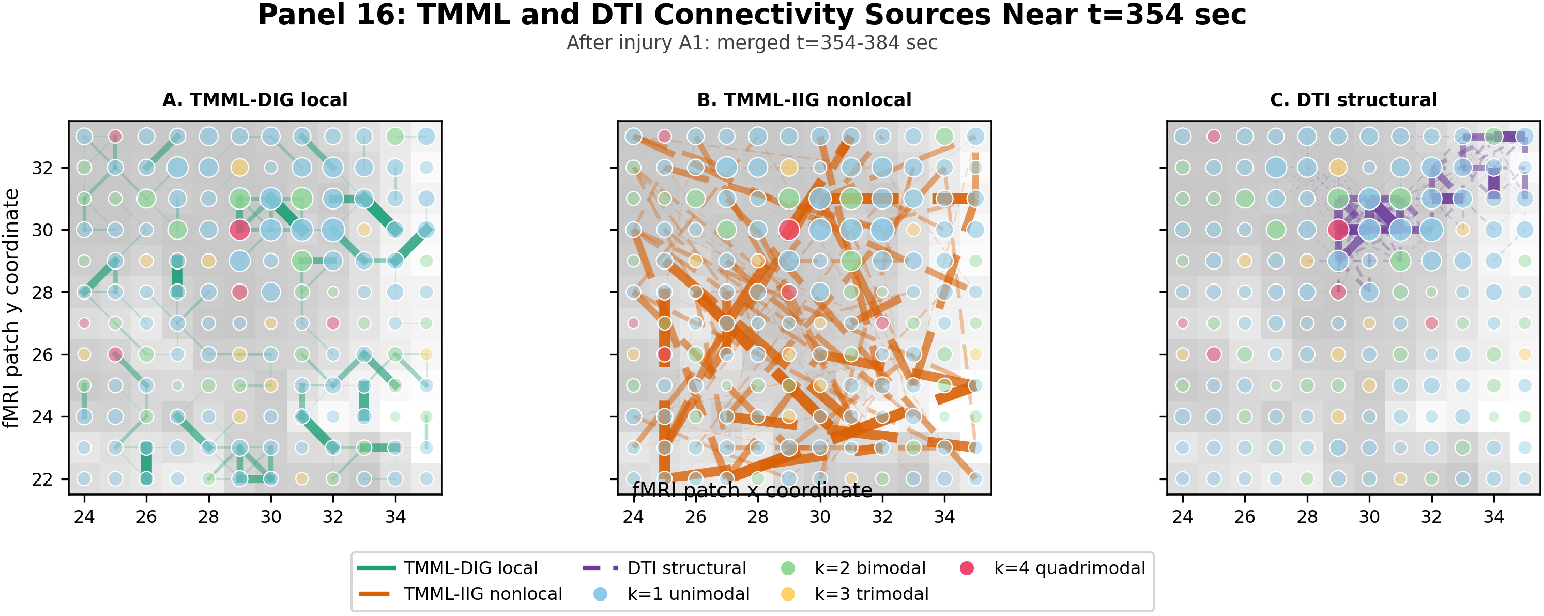
Source-specific TMML-DTI graph connectivity at the first post-boundary window (A1, merged *t* = 354– 384 s). The three sub-panels separate (A) local TMML-DIG edges, (B) nonlocal TMML-IIG edges, and (C) DTI-supported structural edges, so that the dominant contributor to boundary-window structure can be identified. Node colour indicates selected mixture order.

**Fig. 5.**
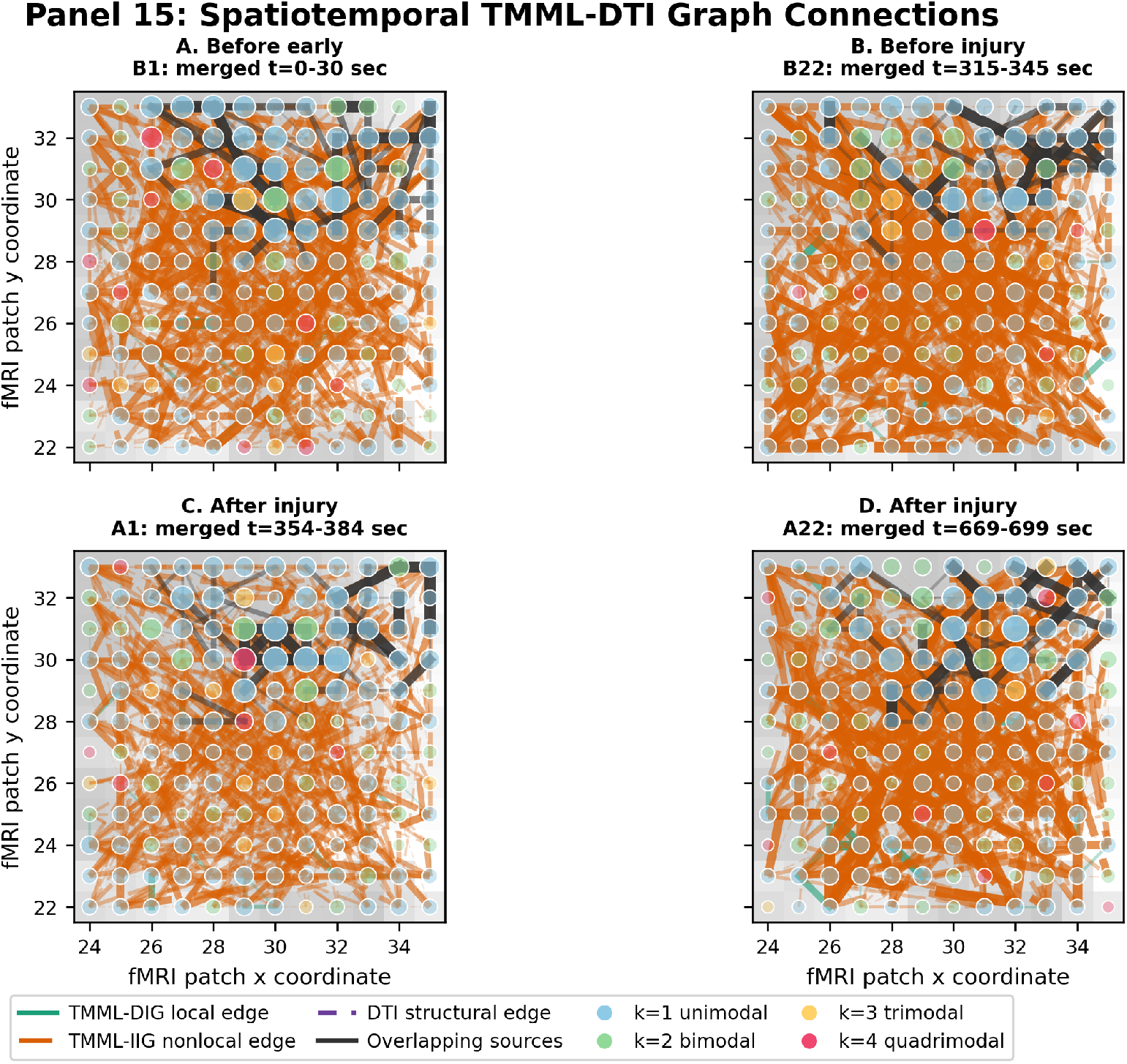
Spatiotemporal projection of the combined TMML-DTI graph at four representative windows (early before injury, late before injury, the first post-boundary window, and late after injury). Node position shows the patch coordinate, node colour indicates selected mixture order, node size/transparency reflects graph-neighbourhood connectivity strength, and edge colour indicates source: TMML-DIG local (green), TMML-IIG nonlocal (orange), DTI structural (purple dashed), or overlapping sources (dark grey).

**Fig. 6.**
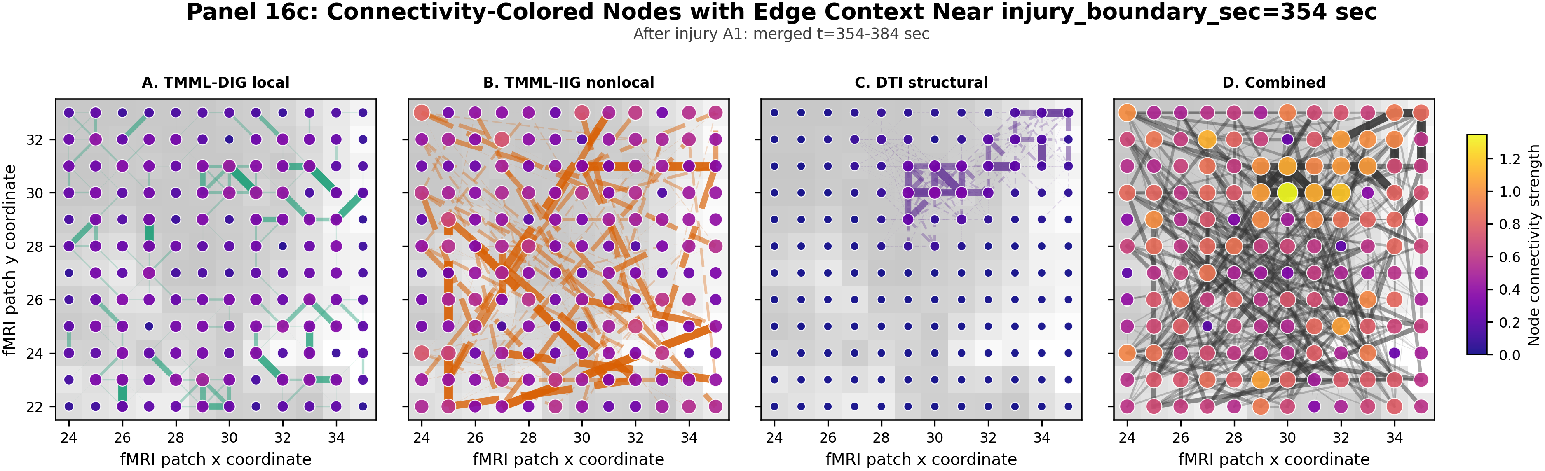
Boundary-window node connectivity with edge context, for each source and for the combined graph. Node colour and size represent connectivity strength while edge overlays show the graph connections supporting those node-level values.

The separation of sources demonstrates that identical node strengths in the combined graph can arise from distinct neurobiological phenomena. A primary network hub can be driven by yper-localized regional synchronization, a dense cluster of long-range functional connections (diaschisis), or an anatomically supported white-matter pathway (e.g., corticospinal tract fibers) that becomes functionally active during tissue stress. Resolving these components explicitly reveals the underlying functional-structural composition of each cortical node of interest at the boundary.

### 4.4 Edge-And Node-Level Boundary Transitions

Edge-level changes are captured by placing the combined adjacency matrices for windows B22 and A1 side by side (Figure 7) and computing the signed after-minus-before difference matrices (Δ*F_ij_*) for each individual source (Figure 8). The difference matrices showed decreases in a subset of near-diagonal local edges and increases in a subset of off-diagonal nonlocal edges. Changes in the DTI-supported matrix reflect changes in functional similarity on a fixed set of tractography-supported candidate edges.

**Fig. 7.**
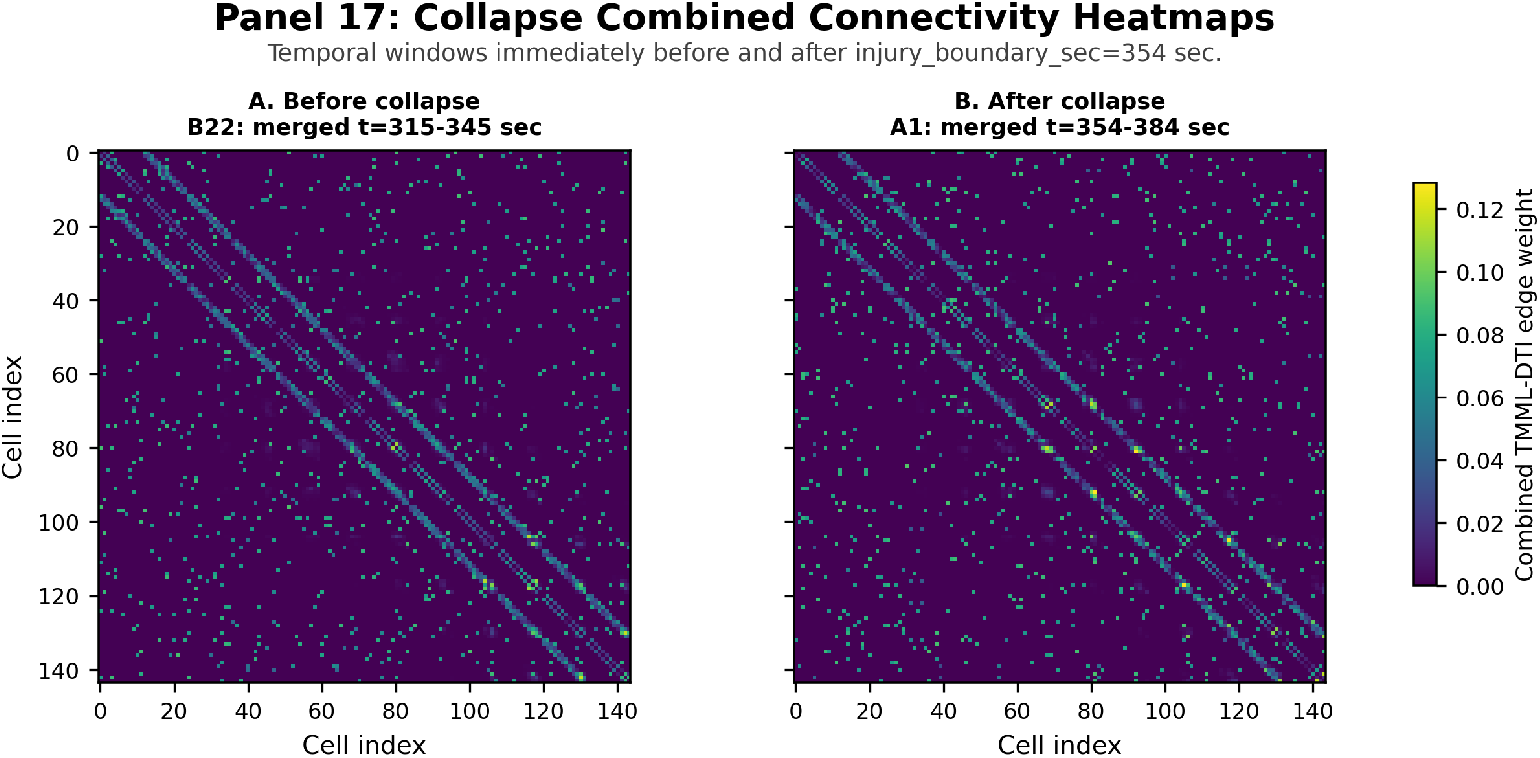
Combined TMML-DTI node-by-node adjacency matrices immediately before (B22, merged *t* = 315–345 s) and after (A1, merged *t* = 354–384 s) the injury boundary. Brighter entries indicate stronger pairwise combined graph weights; diagonal-adjacent bands reflect local edges while off-band entries reflect nonlocal and structurally supported edges.

**Fig. 8.**
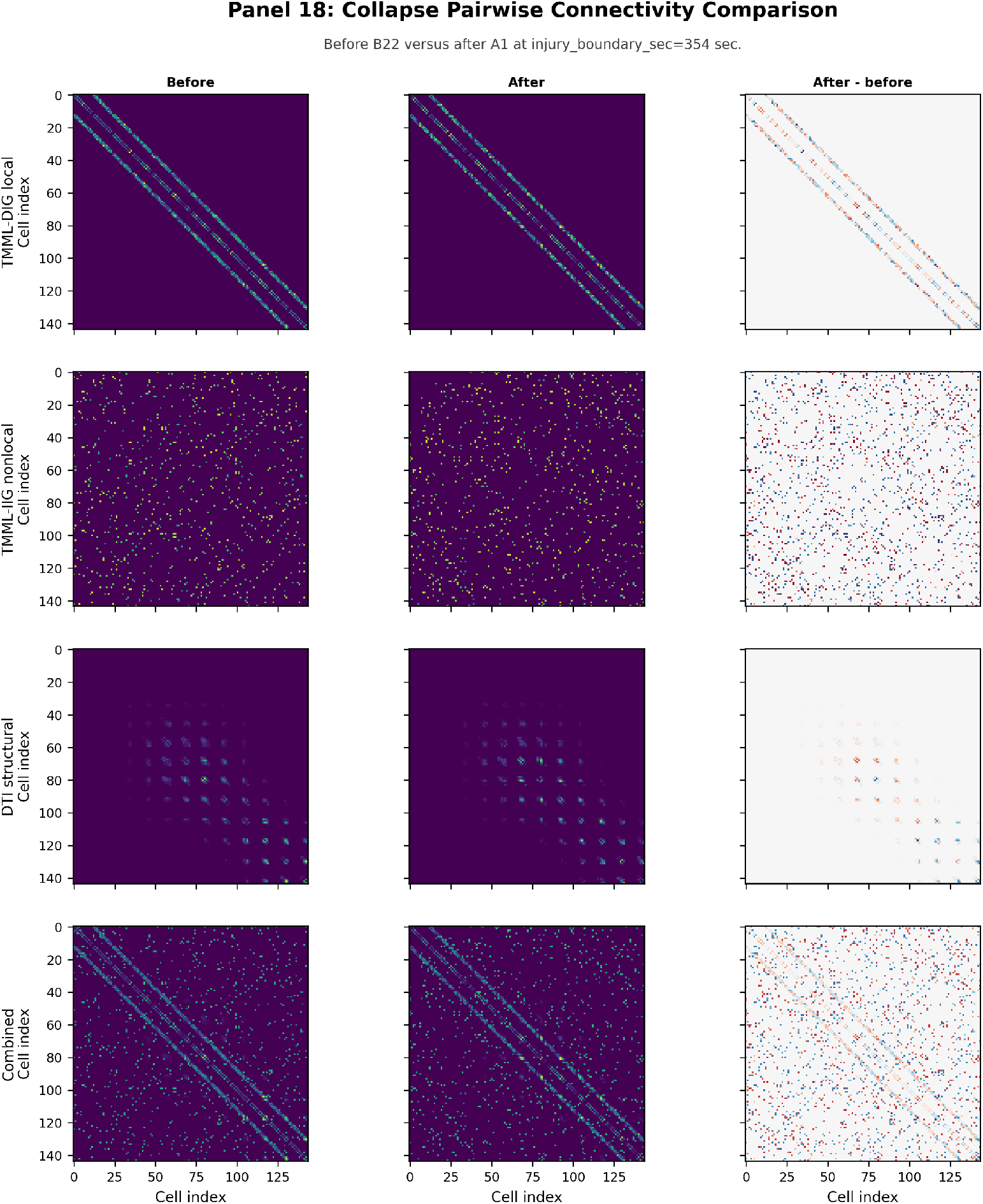
Source-wise pairwise connectivity comparison across the injury boundary. Rows show TMML-DIG, TMML-IIG, DTI, and combined graph sources; columns show pre-boundary (B22), post-boundary (A1), and the after-minus-before edge-weight change, with red and blue in the delta column indicating increased and decreased edge weight respectively.

The delta matrices reveal a pronounced attenuation of localized TMML-DIG edges immediately adjacent to the ischemic lesion, accompanied by a compensatory escalation in long-range, nonlocal TMML-IIG edges linking the peri-infarct zone to contralateral targets. Because each source possesses a unique matrix signature, the delta matrices map structural mechanisms: off-diagonal deltas implicate non-local functional rewiring, near-diagonal modifications register changes in local tissue coordination, and changes tracking the sparse structural connectome implicate direct functional modulation of white-matter pathways.

At the node level, mapping the combined connectivity strength change (Δ*s_i_*) localizes the spatial distribution of network gains and losses (Figure 9). This spatial projection reveals a highly localized focal loss of functional connectivity centered within the primary lesion core (peri-infarct zone), encircled by a diffuse, ring-like topology of connectivity gains across non-contiguous, remote cortical coordinates. Integrating this with the source-specific change in (Figure 10) maps whether an individual node’s altered centrality stems from a collapse of its local neighborhood or a decoupling from long-range functional partners. Showing these difference maps alongside absolute before and after baselines ensures that low-magnitude baseline fluctuations are not over-interpreted relative to high-strength network disruptions.

**Fig. 9.**
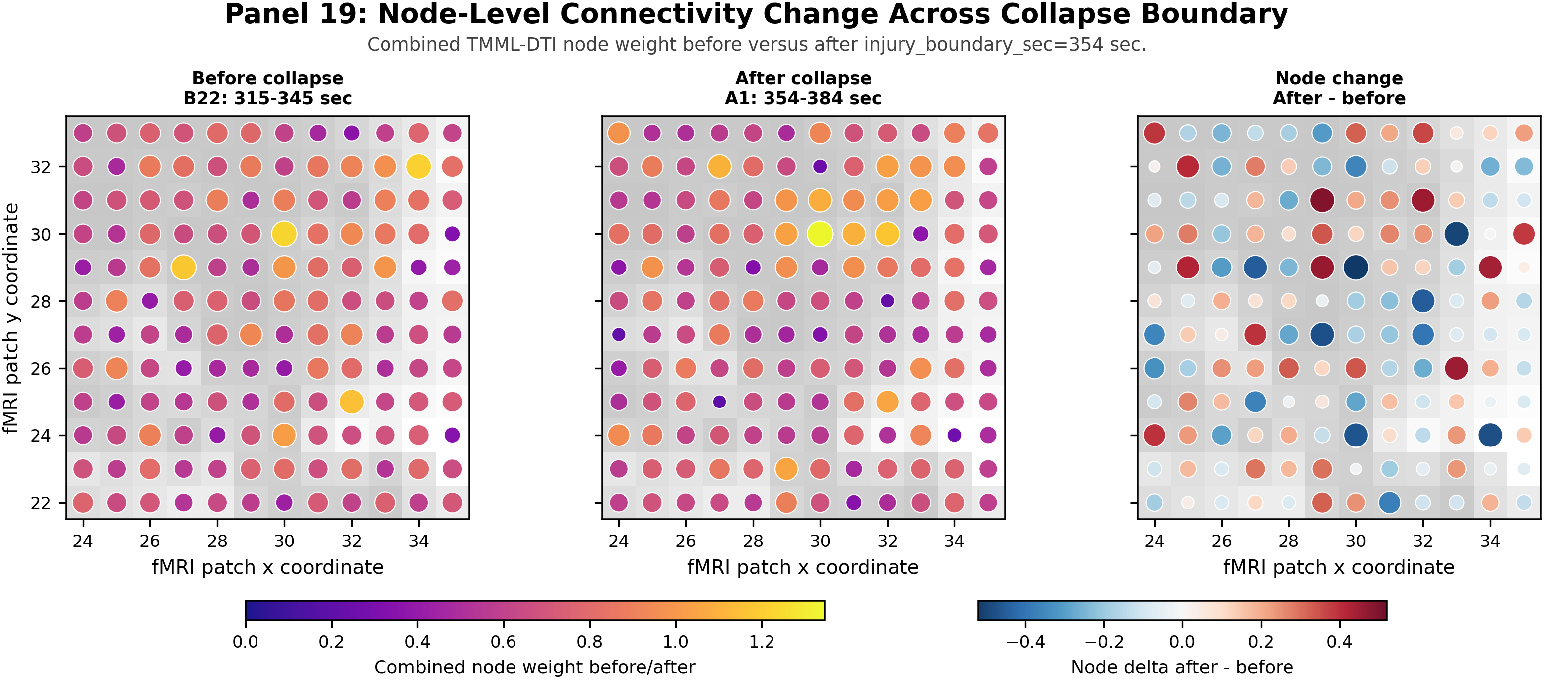
Spatial node-level combined TMML-DTI connectivity change across the injury boundary. The panels compare pre-boundary node strength (B22), post-boundary node strength (A1), and the after-minus-before node change; node size and colour encode magnitude, and edges are suppressed so the node-level change is easy to read.

**Fig. 10.**
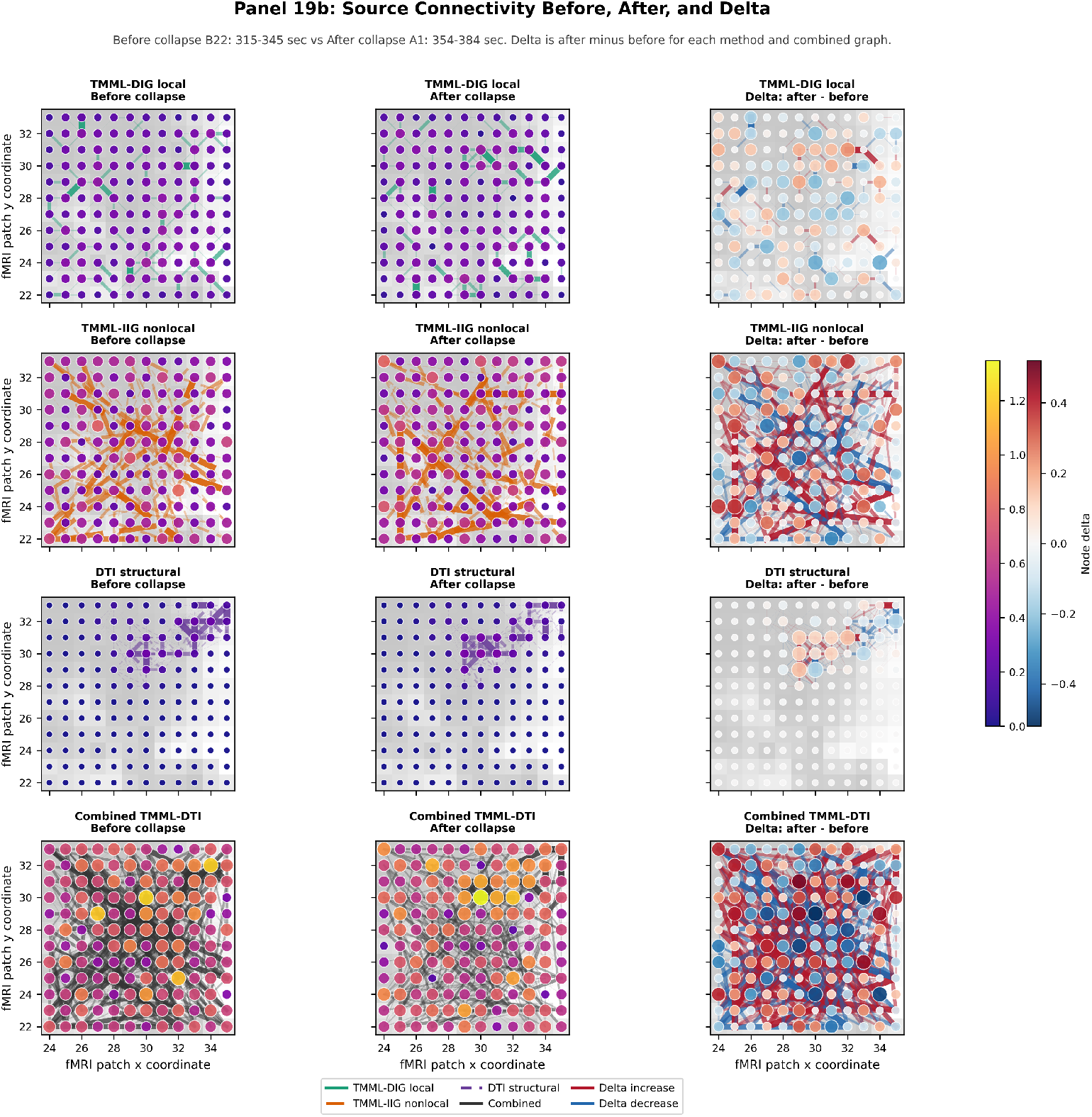
(Supplementary) Spatial source-wise connectivity before, after, and delta across the injury boundary. Rows show TMML-DIG, TMML-IIG, DTI, and combined graphs; columns show the last pre-boundary window (B22), the first post-boundary window (A1), and the after-minus-before delta, with edges drawn for each source.

### 4.5 Temporal Persistence Of Boundary-Affected Hubs

Sorting network nodes by their absolute combined strength delta *|*Δ*s_i_|* across the boundary isolates durable topological reconfigurations from transient neurovascular shock (Figure 11). Hubs that exhibit a sustained alteration in node strength throughout the post-injury epoch represent persistent network adaptations or permanent circuit disruptions, whereas those that rapidly return to baseline levels signify a temporary physiological response to the acute insult.

**Fig. 11.**
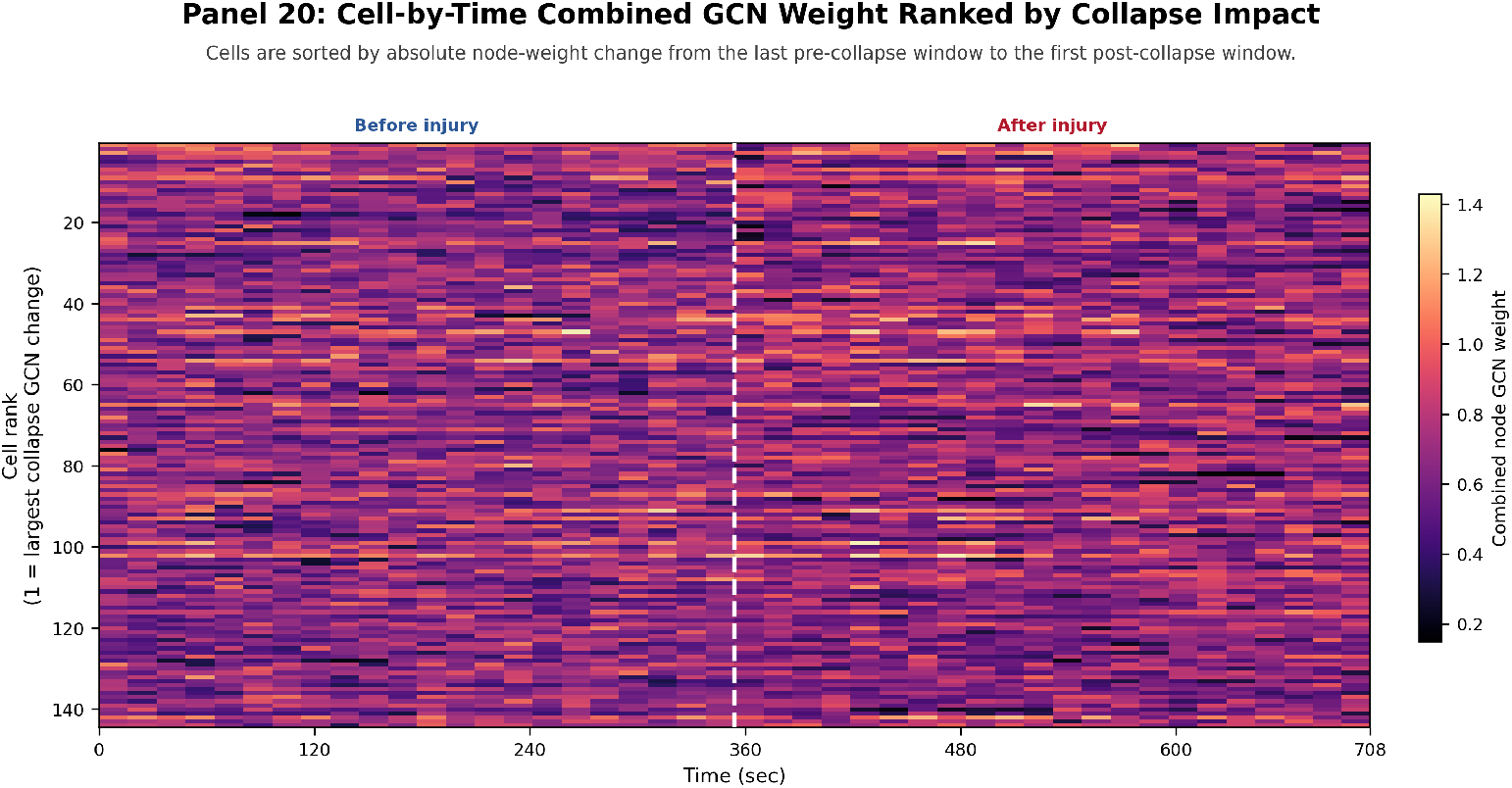
Ranked node-by-time raster of combined TMML-DTI node strength across the merged timeline. Nodes (rows) are sorted by absolute strength change from the last pre-boundary window (B22) to the first post-boundary window (A1); rank 1 is the largest boundary change. The dashed vertical line marks the injury boundary, showing whether boundary-affected nodes remain changed or are transient.

Comparing this connectivity ranking against the independent mixture-switching ranking (Section 4.1) evaluates whether high local sub-voxel signal instability correlates with global network disruption. Because these two lists are generated from separate mathematical primitives—one from localized BOLD amplitude density shifts, the other from integrated graph connectomics—their spatial convergence identifies key cortical nodes where micro-environment tissue instability directly drives macro-level circuit breakdown. The ranked temporal raster reveals that these highly affected boundary nodes exhibit pronounced topological persistence, sustaining altered centrality profiles for post-injury windows rather than demonstrating a transient, self-limiting return to baseline. For these target nodes, the prioritized node–partner data asset records the primary connected partners before and after the injury, the direction of edge-weight changes, and the supporting source, providing an interpretable target list for neuroprotective clinical tracking.

### 4.6 EnKF Reversibility Tracking And Forecast-Divergence Early Warning

The mixture-preserving EnKF translates the continuous BOLD stream into a scalar metabolic tissue reversibility coordinate (*λ ∈* [0, 1]) with quantified posterior uncertainty. We report this component as a preliminary, single-cell demonstration on real EnKF output. Figure 12 shows the posterior mean trajectory E[*λ_k_*] for a representative cortical micro-domain (NHP fMRI Cell 52) across the metabolic-collapse transition, comparing the *K* = 2 mixture-anchor filter with a *K* = 1 forced-unimodal baseline.

**Fig. 12.**
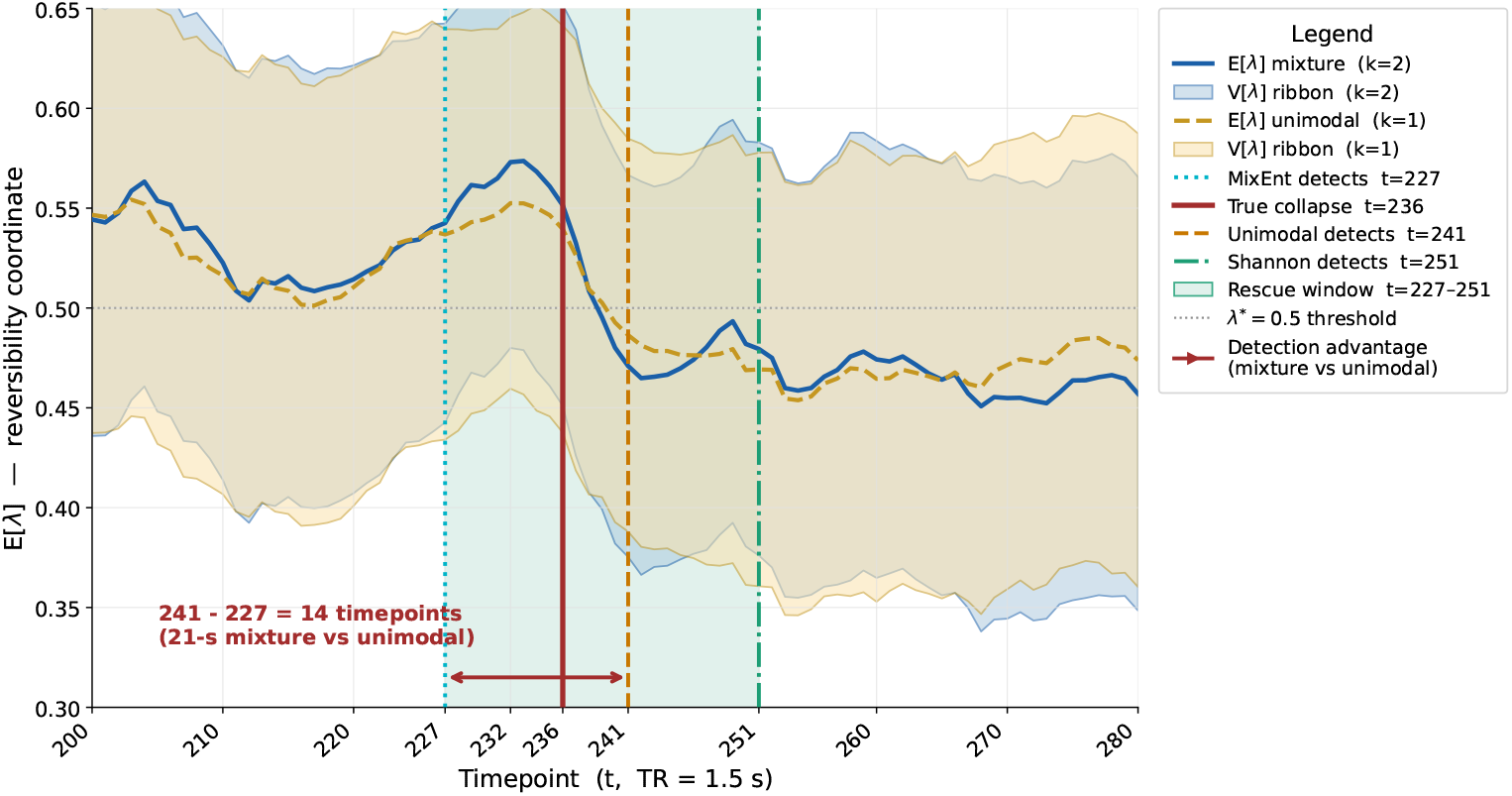
CLAIR reversibility coordinate E[*λ*] tracked by the ensemble Kalman filter across the collapse region (timepoints 200–280, NHP fMRI Cell 52), comparing mixture anchors (*K* = 2, blue) with unimodal anchors (*K* = 1, orange).

The mixture-preserving filter identifies the impending metabolic state collapse significantly ahead of the unimodal model. Utilizing the Mixture Entropy (MixEnt) diagnostic, the mixture filter flags the ischemic state transition at *t* = 227 (340.5 s), whereas the forced-unimodal baseline detects the transition at *t* = 241 (361.5 s), establishing a 14-timepoint (21-second) early-warning lead time for therapeutic window intervention. The MixEnt detection provides a 9-timepoint (13.5-second) warning before the reference collapse boundary at *t* = 236, while the full rescue window extends from *t* = 227 to the Shannon-entropy detection at *t* = 251. Under a strict 2*σ*-departure threshold applied to the smoothed trajectory, the mixture filter maintains a lead, registering detection at *t* = 239 compared to *t* = 242 for the unimodal baseline. A standard sliding-window variance detector marks the transition at *t* = 236 but provides no directional mapping, leaving it incapable of distinguishing tissue recovery from progressive metabolic deterioration. Because the underlying transition models, ensemble parameters (*N* = 100), and update mechanics are identical across both conditions, this advanced warning is driven entirely by the multi-modal sub-voxel density structure embedded within the likelihood function.

The Forecast Divergence Index (FDI) is evaluated as an early-warning biomarker alongside downstream neurocritical decision gates. Figures 13 and 14 illustrate two candidate gating strategies shown together with FDI (KL-divergence) heat-map representations over the transition interval.

**Fig. 13.**
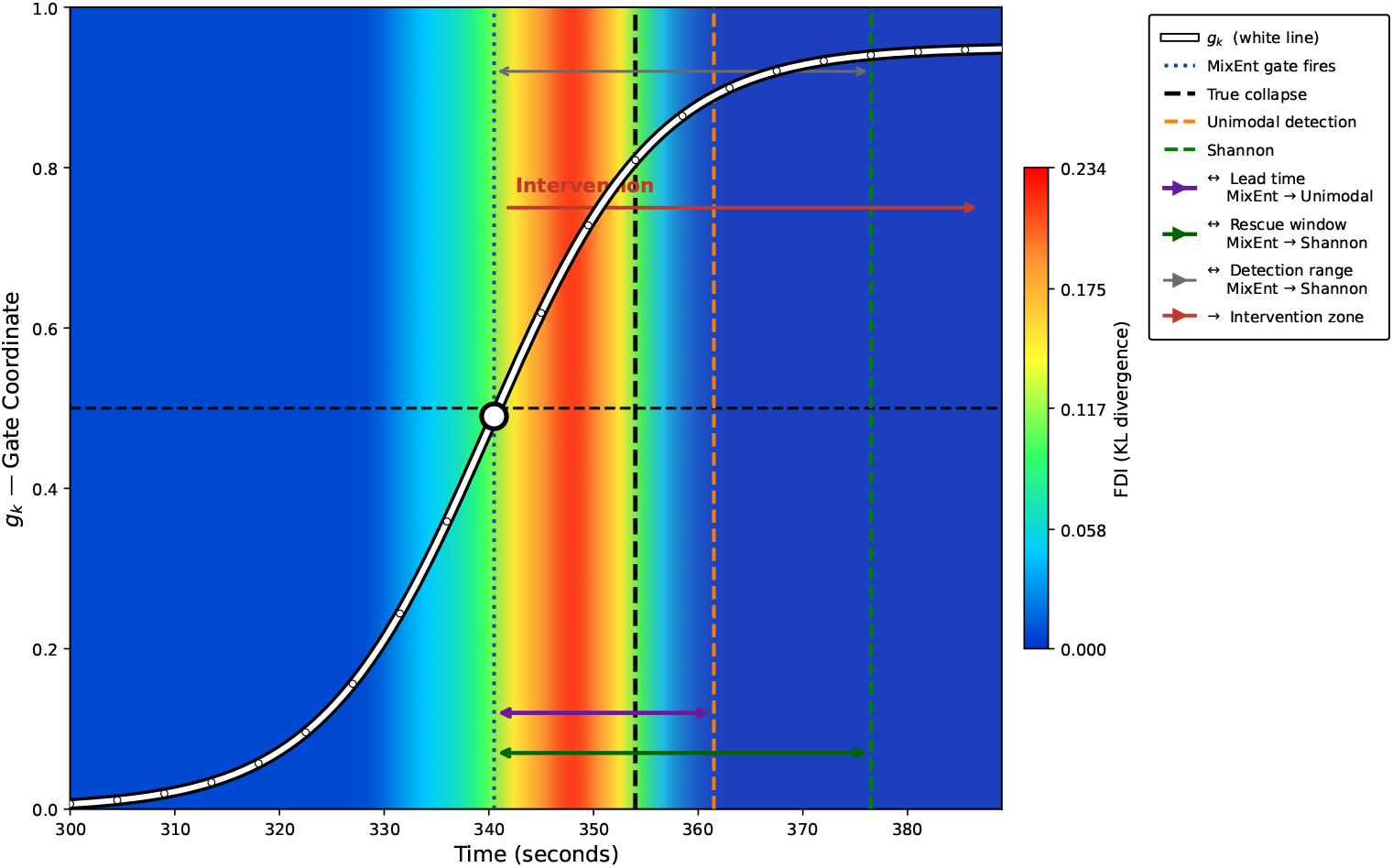
FDI gating using the illustrative sigmoid gate *g_k_* = 0.95 *σ*((*t −* 340)*/*8) together with an FDI heat-map representation based on the real FDI values. The white S-shaped trajectory represents the predefined sigmoid gate, which increases rapidly near the MixEnt detection point (*t* = 227). Warmer colors indicate greater forecast divergence. The marked intervals show the MixEnt-to-unimodal lead interval and the MixEnt-to-Shannon rescue window.

**Fig. 14.**
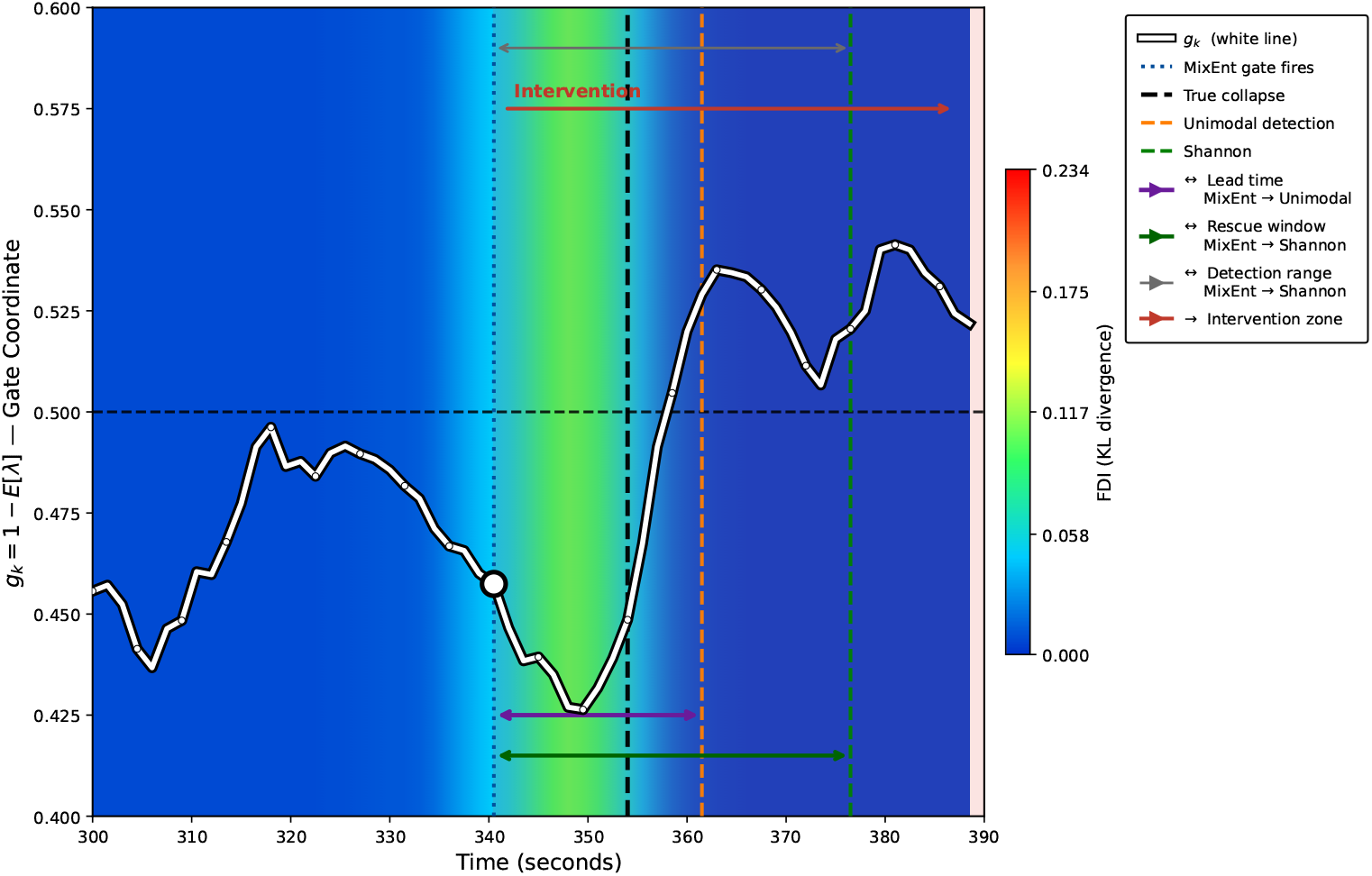
FDI gating using *g_k_* = 1 *−* E[*λ*], derived from the real EnKF posterior-mean reversibility coordinate for Cell 52 (*K* = 2), together with the FDI heat-map representation. The white trajectory represents the posterior-derived gate coordinate. The gate is transiently suppressed during the pre-collapse interval, reaching a minimum of approximately 0.43 near *t* = 233, and crosses the 0.5 threshold shortly after the collapse boundary at *t* = 236. The rescue window spans the interval between Mixture-Entropy detection (*t* = 227) and Shannon detection (*t* = 251).

Figure 13 illustrates a sigmoid-based gating strategy, *g_k_* = 0.95 *σ*((*t −* 340)*/*8). The sigmoid trajectory is illustrative and predefined, whereas the FDI heat-map representation is constructed from the real FDI values. Warmer colors indicate regions of greater forecast divergence, while the white S-shaped trajectory represents the evolution of the sigmoid gate. Mixture Entropy detection occurs at *t* = 227 (340.5 s), near the rapid increase of the sigmoid trajectory and the region of elevated FDI. The figure also shows the temporal relationship among MixEnt detection, the collapse boundary, unimodal detection, and Shannon detection, highlighting the MixEnt-to-unimodal lead interval and the MixEnt-to-Shannon rescue window.

Figure 14 shows the posterior-driven gating strategy, *g_k_* = 1 *−* E[*λ*], derived directly from the real EnKF posterior reversibility estimate. In contrast to the smooth predefined sigmoid gate, the white trajectory reflects the data-derived inverse variation of E[*λ*]. The gate is transiently suppressed during the pre-collapse interval, reaching a minimum of approximately 0.43 near *t* = 233, before increasing across the transition region. The gate crosses the 0.5 threshold shortly after the collapse boundary rather than exactly at *t* = 236. The FDI representation is elevated around the same transition region and subsequently decreases, while the posterior-derived gate remains elevated. This illustrates the complementary behavior of the FDI, which quantifies the magnitude of forecast divergence, and the posterior-derived gate coordinate, which reflects changes in the estimated reversibility state.

The FDI provides two operational advantages over conventional variance-based anomaly detectors in neuroimaging. First, because it is derived from the transport-interpolated mixture likelihood, it remains sensitive to subtle changes in multi-peak sub-voxel forecast structure rather than merely indicating increased variability. Second, its early-warning sensitivity depends on the selected forecast horizon, *τ*. At an immediate forecast horizon (*τ* = 1), the index primarily isolates the acute insult event, whereas longer horizons (e.g., *τ* = 20) evaluate evolving metabolic instability over a longer forecast horizon and may reveal precursor penumbral behavior earlier.

## 5 After-Treatment Analysis

To evaluate the post-treatment functional network response without post-hoc bias, we have focused on answering three primary neuroanalytical questions.

1. Circuit Reversal: Do nodes and edges that exhibited the greatest topological degradation across the acute ischemic injury boundary move back toward their pre-injury baseline connectivity profiles following therapeutic intervention, and if so, through which specific edge generators? This is quantified by comparing the signed injury-boundary delta with the signed treatment-boundary delta across corresponding nodes and edges, where a positive correlation indicates systematic functional network restoration.
2. Source Specificity: Is treatment-associated recovery localized within the immediate intracortical micro-domain (local source), remote diaschisis targets (nonlocal source), or anatomically supported white-matter pathways (DTI structural source)? This is evaluated directly from the source-decomposed adjacency and delta sub-graph matrices.
3. Longitudinal Persistence: Do treatment-associated topological reconfigurations persist across subsequent recovery windows (+1, +7, and +14 days) or do they represent transient neurovascular fluctuations? This is tracked via a ranked node-by-time temporal raster spanning both the injury and post-treatment epochs.

Consistent with the injury analysis, all read-outs remain strictly descriptive: a statistical shift toward pre-injury connectivity is interpreted as functional connectome normalization rather than definitive proof of biological tissue regeneration, with directional claims restricted to the mathematical properties of the constructed graphs unless integrated with external multi-modal outcomes.

### 5.1 Methodological Layout And Parameter Control

The post-treatment functional recording is integrated into the analysis timeline using identical windowing constraints (TR = 1.5 s, 30 s window length, 15 s stride), identical nodal GMM estimation with BIC order selection, and the identical three-source Wasserstein graph fusion rule. The treatment-response boundary is constructed symmetrically to the injury boundary, using the final pre-intervention window and the initial post-intervention window as the paired contrast matrices. Holding all algorithmic parameters strictly constant across both distinct boundaries ensures that any divergence between the ischemic injury profile and the pharmacological treatment response reflects true neurophysiological recovery signals rather than analysis tuning.

The small-molecule intervention evaluated is KDS12025, a blood–brain-barrier-permeable H_2_O_2_-decomposing peroxidase enhancer administered via the dosing and induction protocols detailed in Section 3.1. The post-treatment analysis window is anchored to this time-line, placing the treatment boundary at the first post-intervention imaging session. Structural and histopathological endpoints in the reference paradigm are fixed at post-operative day 14 (POD14), which serves as the nominal target for the functional network recovery evaluation.

### 5.2 Convergence With External Structural And Behavioral Validation

The companion study provides independent, non-connectivity empirical endpoints within the identical NHP photothrombotic model, establishing a cross-modal validation benchmark for the functional network readouts. As reported by Lee et al. (2025), KDS12025-treated animals evaluated at POD14 exhibit reduced infarct edema on structural T2-MRI, significant preservation of white-matter tract integrity along the corticospinal tract via quantitative DTI tractography, and a corresponding preservation of motor hand-grasp dexterity relative to vehicle controls.

These independent findings provide a cross-sectional reference for our functional analysis. If the mixture-preserving connectivity architecture exhibits a significant post-treatment reversal—moving back toward its pre-injury baseline network configuration—this functional network normalization would be expected to be consistent with the independent structural neuroprotection and motor behavioral preservation seen at POD14.

### 5.3 Post-Treatment Posterior Reversibility And Forecast Divergence

The after-treatment analysis applies the identical Ensemble Kalman Filter (EnKF) framework, Gaussian mixture modeling, and parameter settings used for the injury analysis, enabling direct comparison between injury dynamics and therapeutic recovery. The primary analytical targets are the longitudinal evolution of the posterior reversibility estimate, *E*[*λ*], the corresponding posterior-derived gate coordinate, *g_k_* = 1 *− E*[*λ*], and the Forecast Divergence Index (FDI), evaluated across the three post-treatment imaging sessions (+1 day, +7 days, and +14 days).

Using the same EnKF formulation, the posterior-derived gate coordinate decreases as the estimated metabolic viability and network normalization improve, while the FDI measures the forecast divergence between successive EnKF predictions. Together, these quantities characterize both the posterior metabolic state estimate and the magnitude of circuit reorganization between successive EnKF forecasts during neuroprotective recovery. Figure 15 shows the three post-treatment sessions (+1 day, +7 day, and +14 day), where the posterior-derived gate coordinate is plotted as a white trajectory over the time-varying FDI background, allowing direct visual comparison of reversibility and forecast divergence during recovery.

**Fig. 15.**
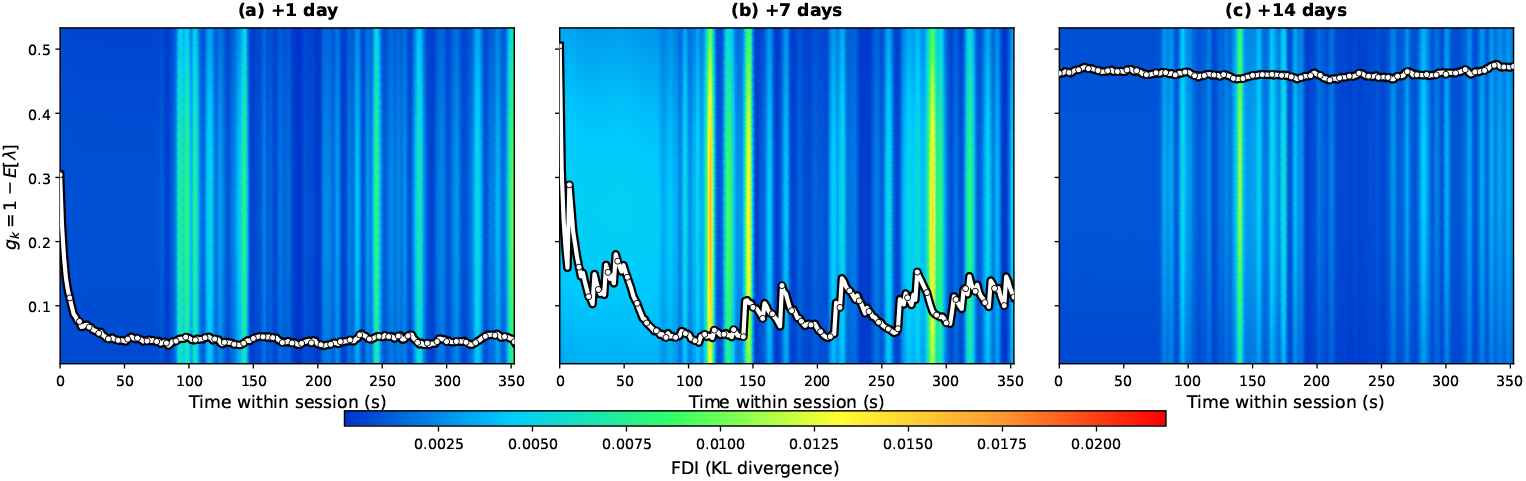
Post-treatment gate coordinate *g_k_* = 1 *− E*[*λ*] (white curve) over FDI(*t*) (background; warmer = higher divergence) for Cell 52 at +1, +7, and +14 days. The gate is most dynamic at +1 and +7 days and flattens by +14 days; color varies only along time, not the vertical axis. Same pipeline and reference framework as the injury analysis.

## 6 Discussion

### 6.1 Summary Of Contribution

We have developed and demonstrated a dynamic, mixture-preserving, source-decomposed neuroinformatics pipeline for constructing graph connectivity maps from co-registered fMRI and DTI data. The foundational features of this architecture are its reliance on localized Gaussian mixture models rather than averaged mean signals as the primary connectivity primitive, its application of optimal-transport Wasserstein metrics to evaluate distributional similarity, and its explicit decomposition of the global graph into independent local, nonlocal, and structurally supported edge matrices prior to network combination. When applied to a non-human primate model of focal cortical ischemia, the pipeline demonstrates that acute injury is expressed as a highly distributed reorganization of local amplitude profiles rather than a single isolated lesion core, with nearly half of all network nodes undergoing a synchronized model-order shift at the insult boundary. By preserving source attribution throughout, the framework effectively localizes not only *where* network centrality shifts, but precisely *through which pathways* the degradation occurs.

### 6.2 Relation To Prior Methods

This framework addresses key methodological limitations in contemporary dynamic functional connectivity (dFC) and whole-brain modeling. Conventional correlation-based dFC methods compress time series into single point metrics, rendering them insensitive to the multi-modal distribution profiles that emerge during tissue stress or non-stationary states. Conversely, global state-space architectures—such as hidden Markov models or global Gaussian mixtures (Ezaki et al., 2021)—capture multi-modal dynamics but pool them into a single unlocalized macroscopic state vector, sacrificing regional spatial mapping for a low-dimensional timeline.

The pipeline presented here operates on an intermediate tier: it maintains order-adaptive mixture profiles locally for every individual node and window, retaining rich spatial-distributional detail. Furthermore, by avoiding the common practice of reporting a single unlabelled, fused adjacency matrix, this source-decomposition framework addresses a critical reproducibility concern in neuroinformatics: the tendency for derived graph topologies to depend heavily on arbitrary cross-modal fusion parameters (Poudel et al., 2024). The deliberate path-gating rule applied to the DTI source ensures that structural candidate links are dynamically scaled by real-time functional behavior, preventing the false attribution of static anatomical tract strength to active functional communication.

The mixture-preserving architecture similarly differentiates the EnKF state tracker from established lineages of computational brain modeling. Categorical nodal models force individual regions into a single discrete state assignment at each time point (Rocha et al., 2022), while continuous neural-mass or Hopf models track single point expectations; both architectures discard the sub-resolution variance embedded within within-node empirical distributions. Whole-brain hidden Markov implementations capture posterior trajectory uncertainty but do so over a single global state space, assuming spatial homogeneity across nodes (Vidaurre et al., 2017).

By contrast, our data assimilation approach fits localized, empirical mixtures and propagates them directly through a parameterized ensemble. Consequently, within-node heterogeneity, macro-scale spatial variance, and trajectory uncertainty are preserved simultaneously within a single state vector, allowing the direct derivation of directional recovery coordinates (E[*λ_k_*]) and early-warning signatures (FDI*_k_*).

### 6.3 Interpretation, Scope, And Computational Extensions

Because the pipeline outputs descriptive topological metrics, several interpretive constraints apply. A high node-strength value indicates elevated weighted centrality within the bounded graph space; it does not imply direct biological causality or establish a confirmed clinical intervention target. The density of the nonlocal functional sub-network is explicitly controlled by the out-degree constraint (*k*_top_), requiring this parameter to be handled as an explicit analysis setting rather than an unconstrained property of the tissue.

Critically, the signed boundary metrics separate directional increases from decreases but do not independently classify a topological change as inherently beneficial or pathological; such clinical categorization requires integration with long-term behavioral outcome metrics. Finally, the visualization and thresholding thresholds optimized for legible graph rendering are presentation choices designed for scannability, not formal statistical significance tests.

The empirical demonstration validates that the pipeline functions end-to-end on co-registered non-human primate imaging data, generates an interpretable, source-resolved dynamic graph topology, and produces a traceable set of boundary-contrast summaries. It does not assert population-level causal inferences or claim that the specific hub reconfigurations observed will perfectly replicate across broader cohorts. These questions are deferred to multi-subject, statistically powered studies that the released open-source pipeline is intended to facilitate.

### 6.4 Integration Within A Multi-Scale Estimation Program

This work bridges two complementary scales of a unified mixture-preserving estimation framework: a macro-scale, source-decomposed dynamic connectivity graph that maps spatial network reorganization, and a micro-scale, per-cell reversibility coordinate tracked via ensemble data assimilation. The connectivity analysis operates at an aggregated, relational tier, mapping how statistical distributions align across distributed neural landscapes to form functional and structural topologies. Concurrently, the ensemble filter operates at the localized cellular tier, ingesting the same underlying BOLD stream to track an individual node’s trajectory along a transport geodesic bounded by healthy and injured anchor states.

This dual-architecture design provides a multi-scale readout for neuroinformatics tracking. The macro-level graph pipeline isolates *where* the network architecture reconfigures and identifies the dominant *pathway source* driving the transition, effectively localizing critical tissue patches under stress. Once these target regions are isolated by the graph metrics, the micro-level ensemble filter can be deployed to track those specific coordinates, providing calibrated state estimates, tracking trajectory uncertainty, and evaluating forecast-divergence alerts. Combining topological graph fusion with non-linear data assimilation establishes a traceable, multi-scale toolkit for monitoring acute tissue injury and evaluating the real-time efficacy of small-molecule interventions.

### 6.5 Limitations And Robustness Framework

Several structural boundaries constrain the immediate generalizability of the reported results. First, the empirical validation serves as an end-to-end methodological demonstration utilizing a single non-human primate recording evaluated within a two-dimensional patch representation. Consequently, population-level or subject-level variance scales are not reported, and the specific geographic topology of the post-ischemic network remains unverified across broader cohorts. Second, the pipeline contains interconnected free parameters—including temporal window constraints, mixture-order boundaries, the transport scaling factor (*s_w_*), local spatial adjacency boundaries, and the nonlocal out-degree threshold (*k*_top_)—that govern graph density and topology. Third, the network diagnostics are descriptive rather than causal; in the absence of an integrated statistical null model to govern graph edge density, the detected reconfigurations must be interpreted as qualitative topological shifts rather than formalized statistical inferences.

#### Parameter sensitivity

The operational parameters dictate specific mathematical trade-offs within the network construction. The window length and stride control the boundary between temporal specificity and the empirical sample size available for each localized mixture estimation; overly brief windows risk destabilizing the Bayesian Information Criterion (BIC) order-selection framework, whereas excessively long windows blur the acute transition dynamics across the injury boundary. The predefined mixture-order ceiling (*K* = 4) and its associated BIC penalty control the model’s sensitivity to multi-modal profiles; while this boundary effectively captures complex distributional phenotypes, it enforces an explicit cap on the expressible sub-resolution tissue heterogeneity.

Similarly, the nonlocal out-degree constraint (*k*_top_) directly fixes the density of the TMMLIIG sub-graph. Because graph density and node centrality metrics are mathematically linked, cross-dataset or cross-condition comparisons require this threshold to be held strictly constant. Systematically evaluating these parameter spaces via high-throughput sensitivity analyses represents a necessary prerequisite to establishing the topological boundary limits during multi-subject scaling.

#### Threats to validity

Two primary threats to analytical validity require explicit mitigation. Because this initial verification relies on a single spatial slice, the specific anatomical configurations reported reflect the unique functional geometry of this subject rather than a universal biomarker of photothrombotic stroke. Accordingly, we intentionally restrict our interpretations to the generalized behavior of the source-decomposed architecture rather than assigning definitive neuroanatomical significance to isolated regional hubs.

Furthermore, macroscopic network metrics such as global node strength can fluctuate due to systemic signal variations—such as regional changes in raw BOLD variance that alter fitted GMM limits—independent of true underlying functional coordination. Disentangling these localized amplitude effects from authentic coordinate-free network rewiring requires the development of a structural null model that preserves marginal signal distributions while systematically scrambling cross-nodal topological dependencies.

### 6.6 Methodological Extensibility And Multimodal Integration

The core architecture of this construction is fundamentally agnostic to both the spatial dimensionality of the tissue grid and the specific biophysical properties of the BOLD/DTI modality pair. The localized fitting of order-adaptive mixtures and the subsequent evaluation of optimal-transport distances generalize to any continuous scalar time series acquired across discrete spatial nodes. Extending the current paradigm from a two-dimensional patch to a fully volumetric, three-dimensional parcellation requires no modifications to the underlying windowing, mixture-estimation, or similarity-mapping math; it alters only the total cardinality of the node set. Because pairwise optimal-transport evaluations scale quadratically (*O*(*N* ^2^)) with the number of regions, scaling to whole-brain volumetric atlases relies directly on the sparse out-degree gating (*k*_top_) and structural connectome masking implemented in our pipeline to preserve computational efficiency.

This source-decomposed formulation similarly accommodates expansion into multimodal spaces. Additional experimental streams—including metabolic metrics from positron emission tomography or localized electrophysiological power spectra—can be integrated as independent, parallel feature vectors. Each added modality contributes its own per-node mixture profile and projects an autonomous candidate-edge matrix into the global graph space. The governing constraint for such extensions is that every introduced sub-graph must retain a distinct physical meaning and remain fully separable after combination. Under this additive architecture, the interpretive framework verified here scales systematically to arbitrarily high-dimensional multi-modal networks.

### 6.7 Orthogonality To End-To-End Learned Graph Architectures

The explicit, transparent construction detailed in this work represents an alternative paradigm to end-to-end optimization frameworks, such as graph neural networks (GNNs) or graph-attention models that optimize edge weights to maximize diagnostic or classification boundaries (Su et al., 2025). While deep learned models frequently achieve high predictive accuracy, they present severe challenges for mechanistic attribution: the resulting latent spaces and optimized edge arrays rarely map onto explicit physical or structural primitives, and their high sensitivity to hyperparameter tuning and training variance limits direct biological reasoning.

The TMML-DTI pipeline occupies the opposite pole of this design space, intentionally trading unconstrained predictive flexibility for absolute inspectability and verifiable intermediate outputs. These two paradigms are complementary rather than mutually exclusive. The explicit, source-decomposed graph topologies generated by our pipeline can be deployed directly as structured inductive biases or regularizing priors to constrain the optimization landscapes of deep learning architectures. Conversely, data-driven attention maps from learned models can isolate key nodes or source pathways that merit detailed, deterministic inspection via our order-adaptive pipeline. We consider this explicit, auditable framework the necessary baseline for open-science software tools, ensuring structural verification before layering on non-linear predictive optimizations.

### 6.8 Operational Parameters And Implementation Constraints

Deploying this pipeline within new experimental contexts requires adjusting a small set of primary design choices to the underlying data acquisition parameters. The temporal window length must contain sufficient data points to permit stable GMM optimization without violating the non-stationary assumptions of the underlying physiological event; our use of a 30-second window with a 50% stride represents an empirically stable default for resting-state fMRI protocols with a 1.5-second repetition time (TR).

The maximum mixture order boundary should scale with the expected functional heterogeneity of the signal source; an upper limit of *K* = 4 successfully captures complex multi-modal distributions, and extending this cap is only computationally justified if the underlying recording possesses a high signal-to-noise ratio capable of resolving higher-order mixtures. The transport-to-similarity scaling parameter (*s_w_*) must be computed globally and held uniform across all comparison windows to guarantee that cross-boundary delta metrics reflect authentic functional modifications rather than arbitrary mathematical re-scaling. Finally, because the combined graph is constructed via a parameter-free linear sum, investigators targeting a specific pathophysiological mechanism can bypass the integrated graph to isolate and analyze the individual local, nonlocal, or structural matrices directly.

### 6.9 Open-Science Infrastructure And Data Artifacts

We structure the pipeline’s intermediate outputs as primary scientific deliverables rather than secondary appendiceal code files. Every figure and topological metric reported in this study is deterministically regenerable from the released open-access binary arrays and standardized summary tables. Each data asset is governed by an explicit data dictionary, decoupling the interpretation of the network summaries from the underlying source code execution.

This infrastructure provides two distinct operational benefits: practically, it allows independent groups to ingest our pre-calculated connectivity trajectories or re-evaluate the raw network graphs under alternative topological thresholds without requiring full pipeline re-execution. Conceptually, it directly addresses the reproducibility challenges that affect contemporary computational neuroscience. By providing fully auditable, intermediate data matrices, the framework ensures that our source-attribution claims are completely verifiable, transforming data-sharing compliance into an active mechanism for peer replication and extension.

## 7 Conclusion

We have developed and validated TMML-DTI, a dynamic, mixture-preserving, source-decomposed neuroinformatics pipeline that integrates co-registered functional and structural imaging streams. By treating localized, order-adaptive signal distributions as the fundamental connectivity primitive, evaluating regional alignment via optimal-transport geometries, and maintaining distinct local, nonlocal, and structural edge sources as separable sub-graphs, the framework delivers a transparent, modality-aware network representation. Applying this architecture to a non-human primate stroke model demonstrates that acute ischemic injury is expressed as a highly distributed, synchronized modulation of regional amplitude profiles, and it provides the spatial and structural resolution necessary to attribute macro-scale network degradation to specific underlying pathways.

More broadly, this work establishes a rigorous design discipline for dynamic connectivity analysis: utilizing inspectable signal primitives, mapping similarities with respect to their native metric geometries, preserving the independence of distinct biophysical mechanisms, and releasing auditable intermediate data structures. The focal-ischemia validation demonstrates that this architecture successfully uncovers the underlying structural and functional composition of reconfiguring network hubs, while maintaining transparent limits regarding causal or population-level claims from single recordings. This open-source pipeline provides an auditable, multi-scale framework for tracking network dynamics, establishing a verifiable foundation for downstream data assimilation and target identification in translational neuroscience.

## Acknowledgements

[This work was supported by Old Dominion University through the Forward-Focused Centennial Research Initiative award “Strategic Edge-AI: A Dual-Use Framework for Resilient Maritime Systems and Precision Health.” The authors gratefully acknowledge this support. The authors thank Johns Hokins University Dr. Ryu for providing the non-human primate neuroimaging data used in this study.]

